# A role for protease activated receptor type 3 (PAR3) in nociception demonstrated through development of a novel peptide agonist

**DOI:** 10.1101/2020.07.08.194373

**Authors:** Juliet Mwirigi, Moeno Kume, Shayne N Hassler, Ayesha Ahmad, Pradipta R. Ray, Changyu Jiang, Alexander Chamessian, Nakleh Mseeh, Breya P Ludwig, Benjamin D. Rivera, Marvin T Nieman, Thomas Van de Ven, Ru-Rong Ji, Gregory Dussor, Scott Boitano, Josef Vagner, Theodore J Price

## Abstract

The protease activated receptor (PAR) family is a group of G-protein coupled receptors (GPCRs) activated by proteolytic cleavage of the extracellular domain. PARs are expressed in a variety of cell types with crucial roles in hemostasis, immune responses, inflammation, and pain. PAR3 is the least researched of the four PARs, with little known about its expression and function. We sought to better understand its potential function in the peripheral sensory nervous system. Mouse single-cell RNA sequencing data demonstrates that PAR3 is widely expressed in dorsal root ganglion (DRG) neurons. Co-expression of PAR3 mRNA with other PARs was identified in various DRG neuron subpopulations, consistent with its proposed role as a coreceptor of other PARs. We developed a lipid tethered PAR3 agonist, C660, that selectively activates PAR3 by eliciting a Ca^2+^ response in DRG and trigeminal (TG) neurons. *In vivo*, C660 induces mechanical hypersensitivity and facial grimacing in WT but not PAR3^-/-^ mice. We characterized other nociceptive phenotypes in PAR3^-/-^ mice and found a loss of hyperalgesic priming in response to IL-6, carrageenan, and a PAR2 agonist, suggesting that PAR3 contributes to long-lasting nociceptor plasticity in some contexts. To examine a potential role of PAR3 in regulating activity of other PARs in sensory neurons, we administered PAR1, PAR2, and PAR4 agonists and assessed mechanical and affective pain behaviors in WT and PAR3^-/-^ mice. We observed that the nociceptive effects of PAR1 agonists were potentiated in the absence of PAR3. Our findings suggest a complex role of PAR3 in the physiology and plasticity of nociceptors.

## INTRODUCTION

Protease activated receptor 3 (PAR3) belongs to the PAR family of G-protein coupled receptors (GPCRs), a group of receptors expressed in many cell types and implicated in a variety of inflammatory pathologies (Cocks and Moffatt, 2000; Steinhoff et al., 2005; Hollenberg et al., 2014; Heuberger and Schuepbach, 2019). Like the other PARs, PAR3 does not have an endogenously present ligand but rather is activated through extracellular cleavage of the N-terminal end via proteases. After proteolytic cleavage, the newly available tethered ligand can bind to the receptor, initiating multiple downstream signaling cascades (Ramachandran and Hollenberg, 2008). In contrast to the other PARs, comparatively little research or drug development efforts have been made for PAR3 since its discovery in the 1990s as a second receptor for thrombin, a protease critical for the coagulation process (Ishihara et al., 1997; Coughlin, 2000; Hamilton and Trejo, 2017). PAR3, encoded by the *F2rl2* gene, is neuronally expressed (Zhu et al., 2005; Chamessian et al., 2018), but its physiological role in sensory neurons in the DRG or TG has not been assessed. PAR3 has been shown to regulate PAR1 signaling in endothelial cells and PAR4 signaling in platelets in response to thrombin (Nakanishi-Matsui et al., 2000; McLaughlin et al., 2007).

Significant roadblocks in PAR3 research have been the lack of specific agonists and skepticism on whether the receptor can signal autonomously. Early research showed that COS-7 cells transfected with human PAR3 stimulated with thrombin were able to trigger robust phosphoinositide signaling (Ishihara et al., 1997). Murine PAR3, on the other hand, was unable to signal on its own in response to thrombin when transfected in COS-7 cells (Nakanishi-Matsui et al., 2000). Studies using agonist peptides based on the tethered ligand sequence of PAR3 (TFRGAP and TFRGAPPNS) have yielded mixed results. TFRGAP elicited a [Ca^2+^]_I_ response in rat astrocytes (Wang et al., 2002) and human smooth muscle cells (Bretschneider et al., 2003). However, it was later observed that TFRGAP induced ERK activation via PAR1 rather than PAR3 in human A-498 carcinoma cells and mouse lung fibroblasts (Kaufmann et al., 2005). Furthermore, studies with PAR3 tethered-ligand sequences have evidenced an inability of PAR3 to self-activate in the absence of other PARs (Ishihara et al., 1997; Hansen et al., 2004). We recently described a lipid tethering approach to profoundly increase the potency of PAR agonist peptides (Flynn et al., 2013). We reasoned that the deployment of this approach for PAR3 could clarify how this receptor might signal in DRG neurons *in vitro* and *in vivo*.

In this study, we had the several aims with the overarching goal of gaining better insight into the potential role of PAR3 in nociception. The first was to characterize PAR3 expression in mouse DRG. We find that *F2rl2* mRNA is widely expressed in nociceptors and overlaps with other PAR-expressing subpopulations. Second, we developed a lipid-tethered peptidomimetic agonist for PAR3 and evaluated its pharmacology *in vivo* and *in vitro*. Finally, we measured both mechanical and affective nociceptive effects of various PAR-mediated and non-PAR-mediated stimuli in PAR3^-/-^ mice. Our findings highlight the role of PAR3 in regulating PAR1- and PAR2-evoked pain behaviors, and hyperalgesic priming.

## RESULTS

### Expression of PAR3 in sensory neurons

Expression of PAR3 has been characterized in megakaryocytes (Colognato et al.), and vascular (Bretschneider et al., 2003; Rosenkranz et al., 2011), and alveolar endothelial cells (McLaughlin et al., 2007). However, not much is known about PAR3 expression in peripheral sensory neurons. To this end, we re-assessed *F2rl2* mRNA expression in mouse DRG single-cell RNA Seq datasets that were generated by Li et al., 2016 (Li et al., 2016). Non-linear embedding and visualization (using t-Distributed Stochastic Neighbor embedding or tSNE) of high-dimensional whole-transcriptome gene expression profiles of individual DRG neurons was performed. It was demonstrated that *F2rl2* mRNA is highly enriched in peptidergic (*Calca*) and non-peptidergic (*P2rx3*) sensory neurons (**Fig. 1A**). Expression of *F2rl2* was also detected in neuronal subpopulations that express *Nppb, Mrgpa3, Mrgprd*, and *Mrgpx1*, all of which are gene markers for distinct populations of pruriceptors (Han et al., 2013; Mishra and Hoon, 2013; Meixiong and Dong, 2017). PAR3 mRNA was also identified in subpopulations of Trpv1-encoding nociceptors, which are crucial for thermal hyperalgesia. A discrete population of *F2rl2* mRNA-expressing sensory neurons was enriched with *F2rl1* (encoding PAR2), which we have recently shown to be crucial for mechanical and affective pain responses in mice (Hassler et al., 2020). Finally, populations of mouse DRG neurons expressing PAR1 (*F2r*), PAR4 (*F2rl3*) and *F2rl2* were also found in these neurons.

**Figure 1.**
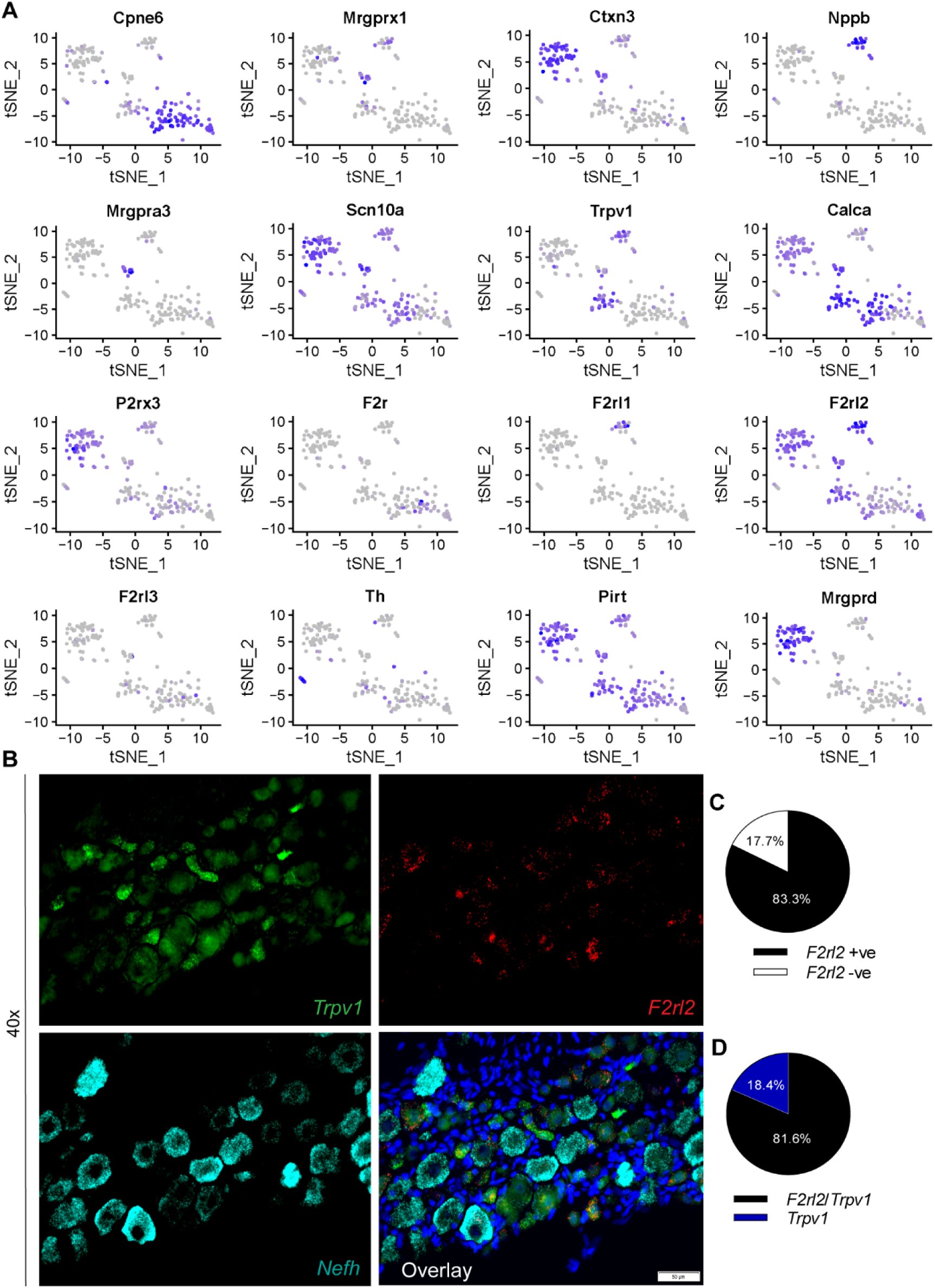
*F2rl2 mRNA* is expressed across sensory neuron populations. t-SNE based visualization (using Seurat) of single cell datasets demonstrate that *F2rl2* mRNA is expressed in a majority of DRG sensory neuron subtypes. *F2rl2* mRNA was detected in populations of sensory neurons co-expressing either *F2r* and *F2rl2* mRNAs that encode PAR1 and PAR2, respectively. Little to no overlap was observed between *F2rl2* and *F2rl3* expressing sensory populations. Of note, *F2rl2* mRNA appears to be broadly distributed among the peptidergic and non-peptidergic subpopulations of sensory neurons, as well as in *Trpv1*+ neurons. A proportion of neurons expressing itch markers *Nppb, Mrgprx1* and *Mrgpra3, Mrgprd* co-expressed *F2rl2 mRNA*. Gene titles are indicated at the top of each t-SNE plot. Color saturation denotes normalized gene expression levels. (B) Representative 40X images of mouse TG neurons labeled with RNAscope *in situ* hybridization for *Trpv1* (green), *F2rl2* (red), and *Nefh* (cyan) mRNAs. Scale bar: 50 μm. Pie charts illustrate that (C) F2rl2 mRNA is present in a majority of TG neurons (83.3%) and, (C) colocalizes with most Trpv1 mRNA expressing neurons (81.6%).

To further extend our studies on PAR3 expression in peripheral sensory neurons, we conducted RNAscope *in situ* hybridization on mouse TG neurons by probing for *Trpv1, F2rl2*, and *Nefh* mRNAs (**Fig. 1B**). Consistent with the findings from mouse DRG single-cell RNA Seq datasets, *F2rl2* mRNA was identified in a majority of TG neurons, approximately 83.3% (**Fig. 1C**). Additionally, most *Trpv1* mRNA-expressing neurons (81.6%) co-expressed F2rl2 mRNA (**Fig. 1D**) thereby confirming the broad expression patterns of PAR3 in peripheral nociceptors.

### Peptidomimetic compound C660 is a selective activator of PAR3

To date, there have been no agonist ligands described that reliably and selectively target PAR3 *in vitro* and *in vivo*. A possible reason is that the receptor does not signal autonomously and, instead, seems to act as an accessory receptor for the activation of other PARs (Hansen et al., 2004). The capability of known PAR3 peptide derivates to activate other PARs further complicates this area. Despite these caveats, we used a synthetic tethered ligand (STL) approach to design selective peptide agonists of PAR3 and evaluated their efficacy *in vitro* using Real Time Cell Analyzer (RTCA) assays (Flynn et al., 2013). A series of lipid-tethered ligands were synthesized by systematic mapping of N-terminal protease-revealed tethered sequences (refer to peptide list in **Supplemental Fig. S1**) and applied to TG neurons at 1 μM to evaluate Ca^2+^ response (**Supplemental Fig. S1**). We used TG neurons because we can generate a larger number of coverslips from fewer animals using TG rather than DRG. C660 (TFRGAPPNSFEEF-pego3-Hdc) elicited the highest Ca^2+^ response at 1 μM when compared to its truncated analogs C661 (GAPPNSFEEF-pego3-Hdc), C662 (TFRGAP-pego3-Hdc), C663 (TFR-pego3-Hdc), C737 (FEEF-pego3-Hdc), and C742 (NSFEEF-pego3-Hdc). Negative control C728 (Ac-pego3-Hdc) and C729 (scrambled C660 peptide PGTEFNFARESFP-pego3-Hdc) were not active.

We then challenged mouse DRG neuronal cultures with 100 nM C660 and found that it elicited a Ca^2+^ response that was comparable to that of cultured TGs (**Fig. 2A**) in terms of the number of cells that responded to treatment. Minimal Ca^2+^ responses were observed in cultured DRG neurons from PAR3^-/-^ mice suggesting that C660 has a specific action at PAR3, at least in the mouse DRG (**Fig. 2B**). In the RTCA assay, HEK293 cells not induced to express PAR3 did not show a response to C660 (**Fig. 2C**). However, in human PAR3-expressing HEK293 cells, C660 induced a physiological response with an EC_50_ of ∼ 900 nM (**Fig. 2D-2E**), again suggesting a specific action of C660 on PAR3. Having established that C660 can induce Ca^2+^ responses in TG and DRG neurons, we sought to test the compound in an independent preparation with a different dependent measure. Spinal cord slices contain intrinsic neurons of the spinal cord and presynaptic terminals of nociceptors from the DRG. In spinal cord slice electrophysiology, C660 increased the frequency, but not amplitude of postsynaptic events in lamina IIo neurons at 10 μM (**Fig. 3A-C**). Tetrodotoxin (TTX) did not influence the effect of C660 on increased frequency of synaptic events in lamina IIo neurons (**Fig. 3D-F**). Because the frequency of these synaptic events is determined by presynaptic neurotransmitter release and the amplitude is due to postsynaptic receptor density, this finding supports the conclusion that C660 acts on presynaptic PAR3 expressed by DRG neurons to induce neurotransmitter release onto lamina II neurons in the dorsal horn.

**Figure 2.**
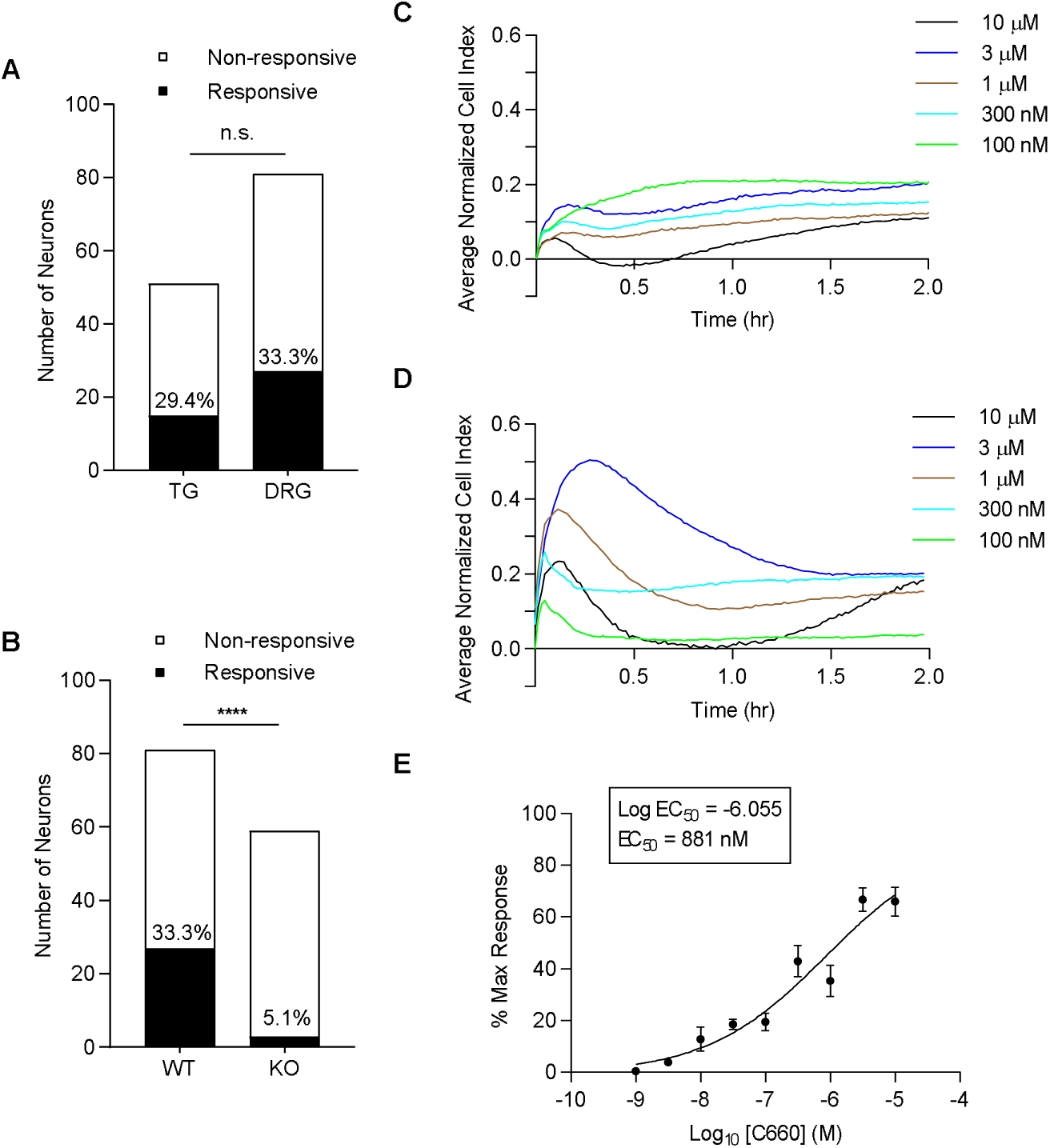
Peptidomimetic agonist C660 elicits responses specific to PAR3-expressing cells. (Panel A and B) WT and PAR3^-/-^ TG or DRG cultures were treated with PAR3 agonist peptide, C660 (100nM), after which the neuronal responses were evaluated based on a 10% ratiometric increase in 340nm/380nm. Only cells with a response to KCl (50 mM) were considered for analysis. Responses in calcium imaging are shown as contingency graphs. (A) C660 (100nM) evokes a comparable Ca^2+^ response in TG (29.4%) and DRG (33.3%) neurons (n= 51 and 81, respectively). (B) C660-evoked Ca^2+^ responses are specific PAR3-expressing DRG neurons as demonstrated by the minimal Ca^2+^ responses from cultured PAR3^-/-^ DRG neurons (n = 59). Fisher’s exact test, ****p<0.0001, n represents number of neurons imaged. Panels (C-E) C660 elicits similar response and selectivity in PAR3-induced HEK293 cells. Cell Index (CI) over time (hr) for non-induced (C) and PAR3-induced (D) HEK293 cells during and after treatment with varying concentrations of C660. CI was recorded every minute for four hours (2 hours shown), with individual traces representing the average of a quadruplicate from a single plate. Changes in peak values were used to construct (E) a concentration response curve, with a calculated EC_50_ of 881 nM for human PAR3. Data points represent mean ± SEM, with n = 16, 16, 16, 15, 16, 3, 8, 6, and 1, respectively, in descending concentration, where n represents each peak difference for a given concentration.

**Figure 3.**
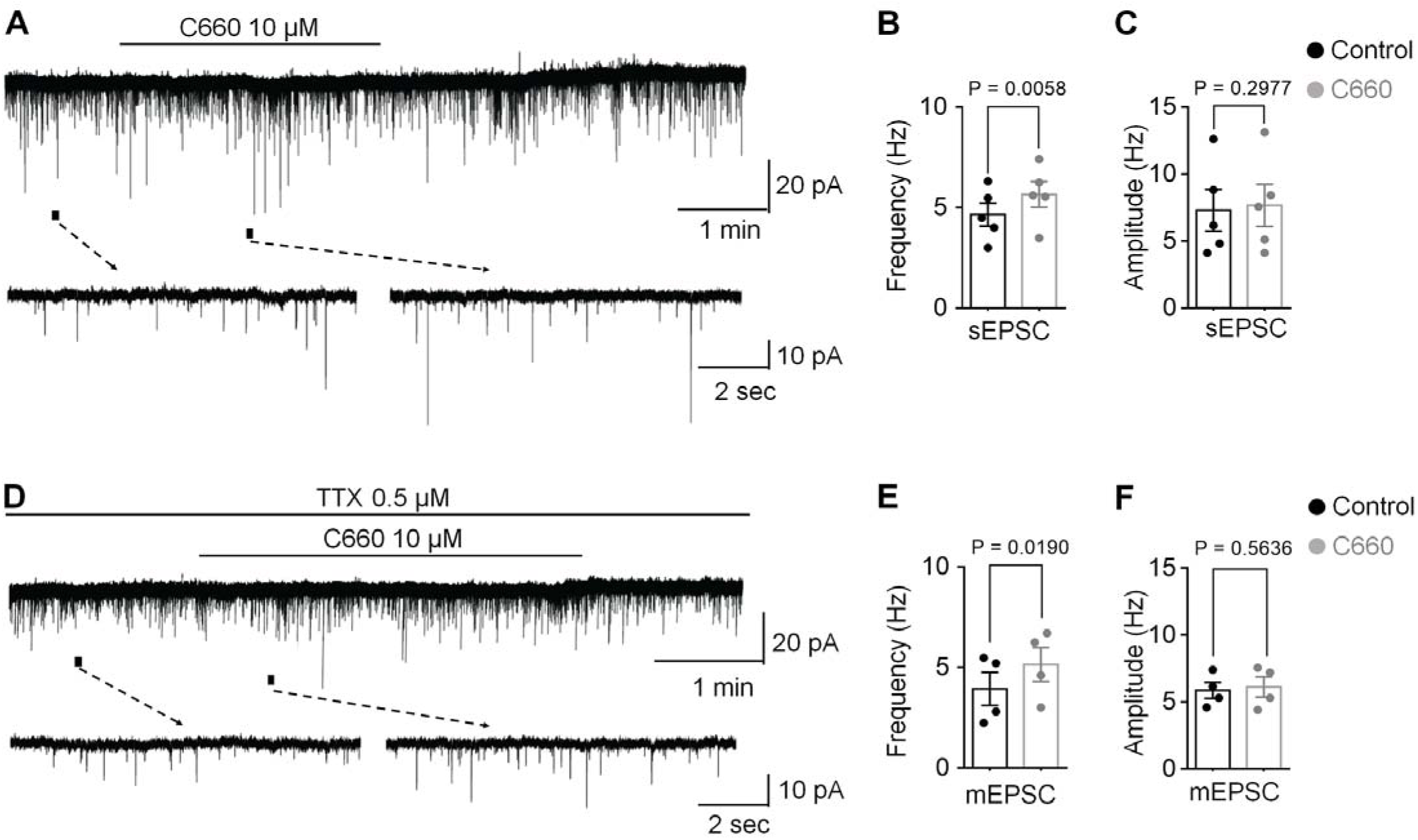
Peptidomimetic compound C660 acts presynaptically to increase dorsal horn excitatory synaptic transmission in spinal cord slices. Whole-cell voltage clamp recordings of lamina IIo neurons in lumbar segments of the spinal cord. (A) Representative traces of spontaneous excitatory postsynaptic currents (sEPSCs) before and after application of C660 (10 μM). Lower: Enlarged traces of events for a period indicated by short bars. sEPSCs were recorded at a holding potential (V_H_) of -70 mV in the presence of 10 μM picrotoxin and 2 μM strychnine. (B, C) sEPSC frequency (B) and sEPSC amplitude (C). Treatment of lumbar spinal cord slices with 10 μM C660 significantly increased the frequency but not amplitude of recorded sEPSCs. n = 5 neurons/group. (D) Representative traces of miniature excitatory postsynaptic currents (mEPSCs) in lamina IIo neurons recorded in the presence of 10 μM picrotoxin, 2 μM strychnine and 0.5 μM tetrodotoxin (TTX). Lower: enlarged traces of events for a period indicated by short bars. (E, F) mEPSC frequency (E) and mEPSC amplitude (F). C660 treatment (10 μM) increased the frequency of mEPSCs with no change in amplitude (n = 4 neurons/group). Numerical data is represented as mean ± SEM. Statistical significance was determined as *P* < 0.05 using paired Student’s *t* test. In all experiments, n refers to the number of the neurons studied. Only one neuron was recorded in each slice.

The involvement of PAR3 in modulating pain behaviors is not well understood due to the scarcity of pharmacological tools that specifically target PAR3 *in vivo*. Therefore, having confirmed the selectivity of C660 for PAR3 using *in vitro* assays, we proceeded to evaluate mechanical and affective pain responses *in vivo*. Mice were injected with 30 pmol of C660 (dosage was estimated from the EC_50_) on the hind paw after recording baseline (BL) measures. von Frey and grimace tests were performed at 1, 3, 5, 24, and 48 hours post-injection. Consistent with our *in vitro* findings, C660 evoked prolonged mechanical hypersensitivity and hyperalgesic priming in wildtype (WT) mice (**Fig. 4A -D**). On the other hand, in PAR3^-/-^ mice, C660 had little acute effect and the magnitude of the hyperalgesic priming effect was greatly reduced in these mice (**Fig. 4A - D**). Facial grimacing following C660 injection was noted in WT mice, although changes were transient (**Fig. 4B**). This suggests that PAR3 activation causes mechanical hypersensitivity that is prolonged and an affective pain response that is relatively brief. Our results show that C660 is a specific agonist of PAR3 *in vitro* with efficacy and selectivity *in vivo*.

**Figure 4.**
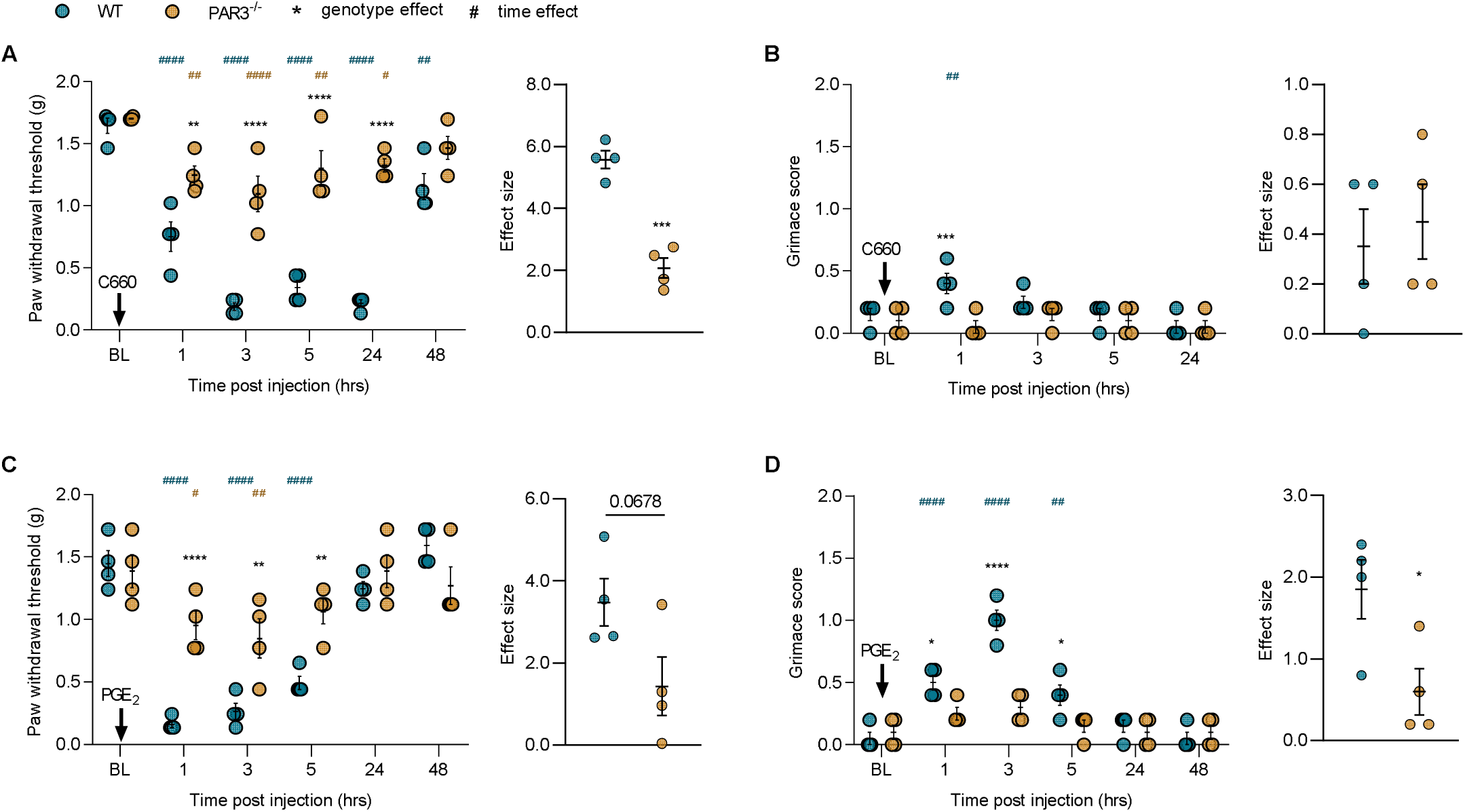
C660 acts on PAR3 to induce hyperalgesia and hyperalgesic priming. WT and PAR3^-/-^ mice were injected with C660 (30 pmol) intraplantarly after recording baseline (BL) measures for paw withdrawal threshold and grimacing. Mechanical and affective measures of pain were subsequently scored at 1, 3, 5, 24, and 48 hours post injection. Effect sizes were calculated to compare the cumulative differences from baseline between the WT and PAR3^-/-^ groups over a duration of 48 hrs post injection. (A) PAR3 agonist, C660, significantly reduced paw withdrawal thresholds in the WT but not PAR3^-/-^ group, with effects lasting up to 24 hrs post injection (n = 4/group). (B) A transient increase in facial grimacing was observed with the WT cohort at the 1 hr time point. However, the overall effect size over a 24 hr period did not differ significantly between groups (n = 4/group). To assess the effects of C660 on hyperalgesic priming, PGE_2_ (100 ng) was applied to the hind paw of WT and PAR3^-/-^ mice 14 days after initial stimulation with C660 (30 pmol) (Panel C and D). (C) Significant genotypic differences in hyperalgesic priming were observed at the 1, 3, and 5 hr timepoints and were resolved after 24 hrs. Cumulatively, PGE_2_ hyperalgesia was partially attenuated in PAR3^-/-^ cohort (n = 4/group). (D) PGE_2_ produced a robust affective response in WT but not PAR3^-/-^ mice. Data is expressed as mean ± SEM. Two-way ANOVA with Bonferroni’s multiple comparisons (Panel A-D) *p<0.05, **p<0.01, ****p<0.0001. Unpaired t-test *p<0.05, ***p<0.001. Stars show significant differences between treatments or genotypes. Hashtags show differences by time, from baseline.

### Knockout of PAR3 potentiates pain responses to other PAR agonists

Considering that PAR3 is widely regarded as a co-receptor for other PARs, much focus has been drawn to its interactions with other PARs. In endothelial cells, for example, PAR3 is thought to form constitutive heterodimers with PAR1 that favor distinct signaling pathways from PAR1/PAR1 homodimer signaling (McLaughlin et al., 2007). However, the nature of these interactions in nociceptors and the pain behaviors they might elicit as a consequence, are not well understood. We surmised that a knockdown of the non-signaling receptor PAR3 would impact mechanical and affective pain responses to other selective PAR agonists. Interestingly, we observed that PAR1 agonists, thrombin (10 units, i.pl) and TFFLLR-NH_2_ (100 μg, i.pl) induced mechanical hypersensitivity in both WT and PAR3^-/-^, but these responses were significantly more robust and prolonged in PAR3^-/-^ mice. Additionally, facial grimacing was noted in the PAR3^-/-^ mice up to 5 hours post injection with either thrombin or TFFLLR-NH_2_ (**Fig. 5A-B**).

**Figure 5.**
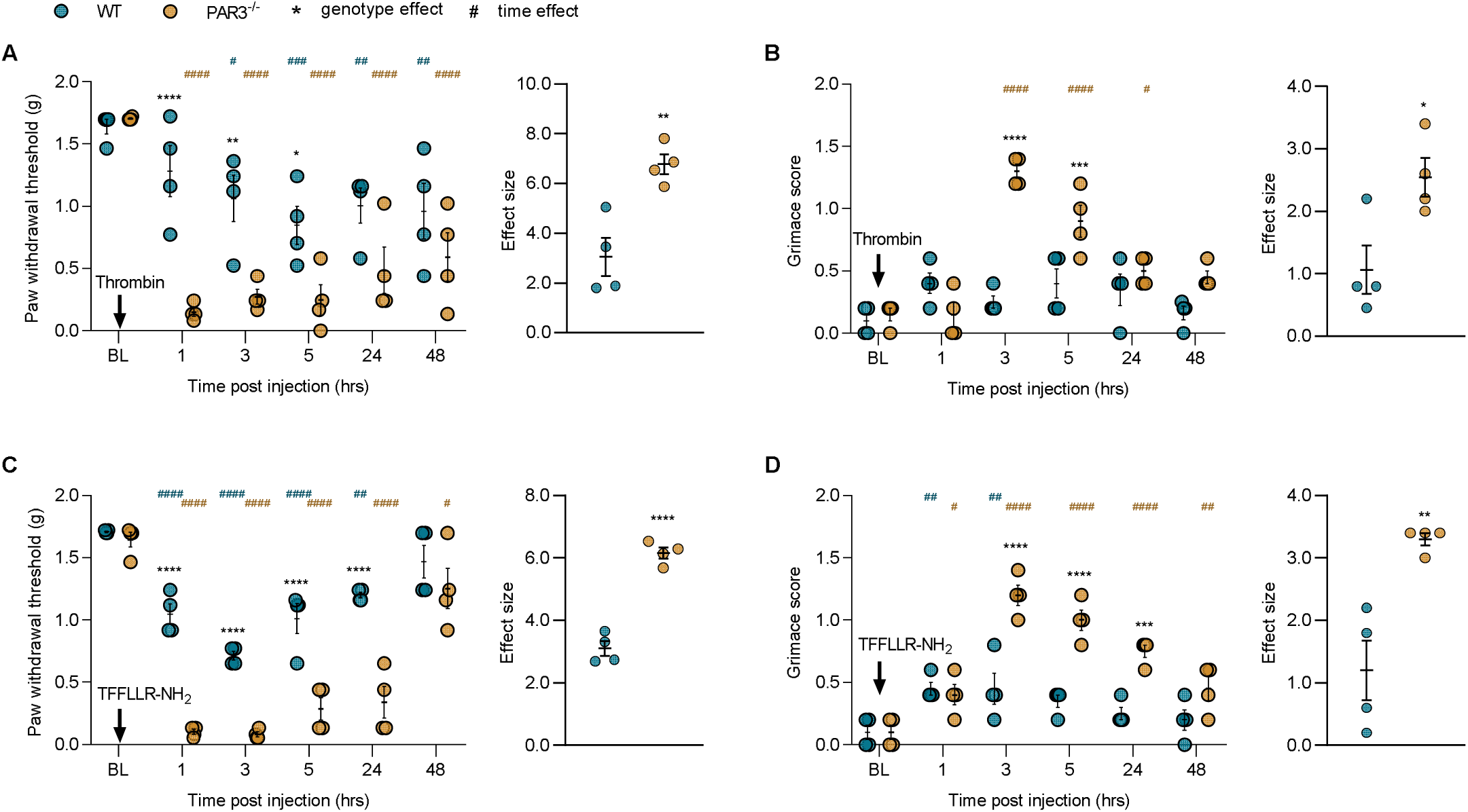
Pronociceptive effects of PAR1 agonists are increased and prolonged in PAR3^-/-^ mice. Thrombin or peptide PAR1 agonist, TFFLLR-NH_2_, was administered intraplantarly into the hind paws of WT and PAR3^-/-^ mice. Baseline (BL) recordings of mechanical thresholds and grimace scores were noted before administering either thrombin (10 units, i.pl) or TFFLLR-NH_2_ (100 μg, i.pl). Thereafter, paw withdrawal thresholds and grimace scores were assessed 1, 3, 5, 24, and 48 hours post injection. (A) Thrombin (10 units, i.pl) induced lasting mechanical hypersensitivity in both WT and PAR3^-/-^ mice. However, the response to thrombin was significantly greater in PAR3^-/-^ mice at 1, 3, and 5 hours post injection (n = 4/group). (B) Thrombin (10 units, i.pl) significantly increased facial grimacing in PAR3^-/-^ but not WT mice (n = 4/group). (C) Intraplantar injections of TFFLLR-NH_2_ (100 μg) significantly reduced mechanical withdrawal thresholds in WT and PAR3^-/-^ mice but was more marked in the latter group. Cumulatively, the genotypic differences over the duration of 48 hours post injection were significant (n = 4/group). (D) TFFLLR-NH_2_ (100 μg, i.pl) induced prolonged facial grimacing in PAR3^-/-^ but not WT mice (n = 4/group). Data is expressed as mean ± SEM. Two-way ANOVA with Bonferroni’s multiple comparisons (Panel A-D) *p<0.05, **p<0.01, ***p<0.001, ****p<0.0001. Unpaired t-test *p<0.05, **p<0.01, ****p<0.0001. Stars show significant differences between treatments or genotypes. Hashtags show differences by time, from baseline.

We next evaluated pain responses after injecting the PAR2 agonist, 2AT (30 pmol) into the hind paw of WT, and PAR3^-/-^ mice. 2AT evoked mechanical hypersensitivity and facial grimacing in both WT and PAR3^-/-^ mice without any effect of genotype (**Fig. 6A-B**). We also assessed hyperalgesic priming in these mice because our previous work demonstrated that PAR2 activation is sufficient to induce priming (Tillu et al., 2015). Unexpectedly, when we challenged these mice with PGE_2_ injection into the previously stimulated hindpaw, we observed a profound deficit in priming in the PAR3^-/-^ mice in the von Frey assay (**Fig. 6C-D**). To test whether similar effects would be observed in the TG system, we injected 2AT (30 pmol) onto the dura of WT and PAR3^-/-^ mice to assess migraine pain-like behaviors (Burgos-Vega et al., 2018; Hassler et al., 2018). 2AT administered supradurally caused robust periorbital mechanical hypersensitivity and grimacing in both genotypes (**Supplementary Fig. S3A-B**). When we assessed priming by applying pH 7.0 solution to the dura (Burgos-Vega et al., 2018) we again did not see any difference between genotypes (**Supplementary Fig. S3C-D**).

**Figure 6.**
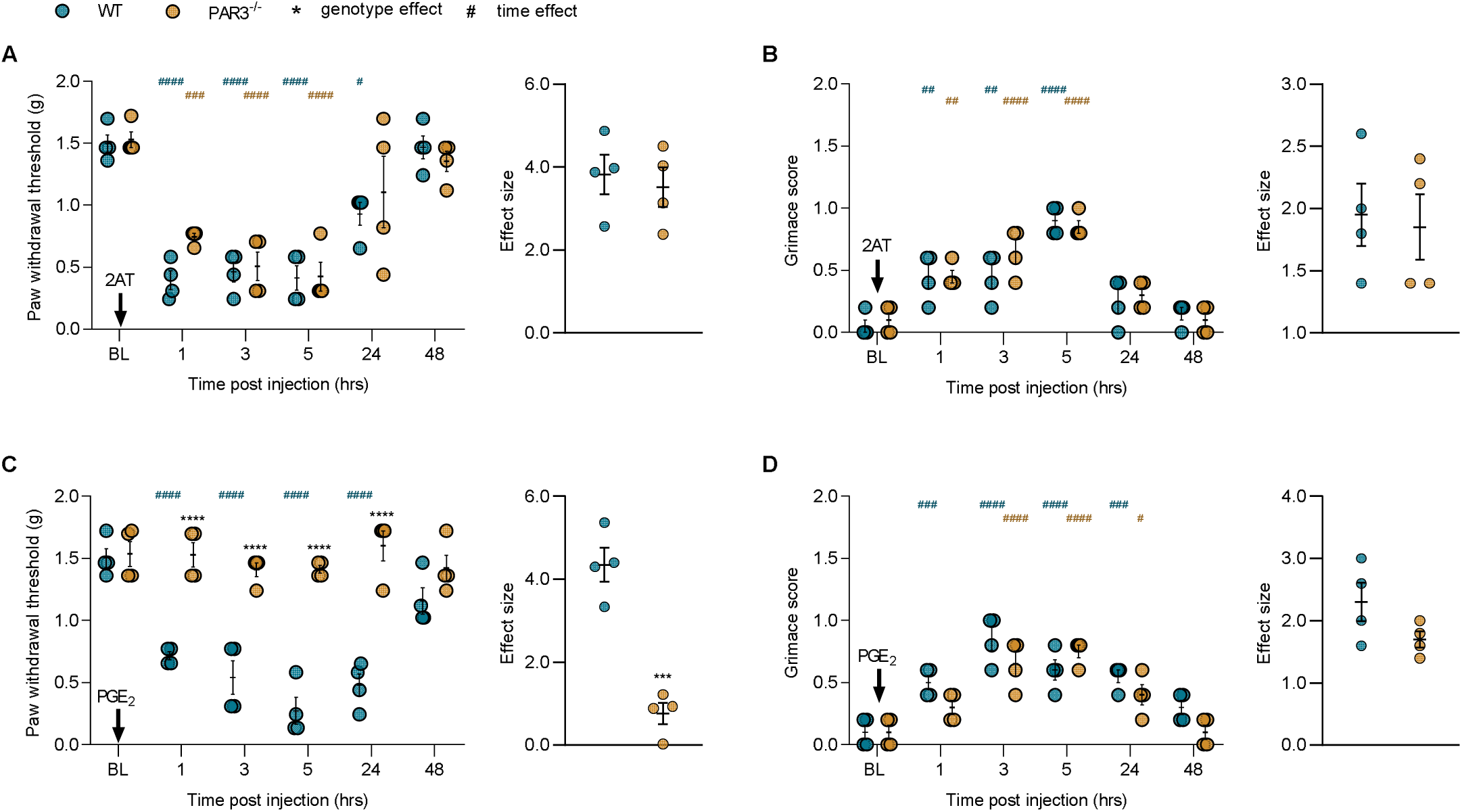
PAR2 agonist, 2AT, induces mechanical hypersensitivity and facial grimacing but not hyperalgesic priming in PAR3 deficient mice. WT and PAR3^-/-^ mice were injected with 2AT (30 pmol) into their hind paws and then mechanical and affective measures of pain were recorded up to 48 hours. Effect sizes were calculated to compare the cumulative differences from BL over a 48 hr duration for the WT and PAR3^-/-^ groups. 2AT injected into the hind paw significantly increased (A) mechanical hypersensitivity (B) and facial grimacing (n = 4/group) in both WT and PAR3^-/-^ mice, with effects lasting up to 5 hours. (Panel C and D) WT and PAR3^-/-^ mice pretreated with 2AT (30 pmol), received an i.pl injection of PGE_2_ (100 ng) in the hind paw 14 d later. Mechanical and affective measures of pain were assessed via von Frey testing and mouse grimace scale respectively (n = 4/group). (C) Mechanical hyperalgesia after PGE_2_ (100 ng) was robust in the WT group only. The time effect for the WT cohort was statistically significant up to 24 hours post PGE_2_ injection. Unpaired t-test of the effect sizes reveal a significant genotype difference between WT and PAR3^-/-^ groups (n = 4/group). (D) Facial grimacing was increased in both groups after PGE_2_ (100 ng) injection (n = 4/group). Data is expressed as mean ± SEM. Two-way ANOVA with Bonferroni’s multiple comparisons (Panel A-D) *p<0.05, **p<0.01, ***p<0.001, ****p<0.0001. Unpaired t-test ***p<0.001. Stars show significant differences between treatments or genotypes. Hashtags show differences by time, from baseline.

The mast cell degranulating compound 48/80 produces pain that is mediated, at least in part, by mast cell tryptase action on PAR2 (Bunnett, 2006). We tested the effect of 48/80 injection into the paw in WT and PAR3^-/-^ mice (**Supplementary Fig. S4A-B**). The effect of 48/80 on mechanical hypersensitivity was blunted in PAR3^-/-^ mice compared to WT mice and there was little grimacing effect observed in this experiment. Therefore, PAR3 does not seem to regulate PAR2-mediated pain responses in response to direct agonist stimulation of the receptor in the DRG or TG regions but there is a contribution of PAR3 to hyperalgesic priming, specifically in the DRG region. PAR3 may play a more significant role in pain responses when PAR2 is activated by endogenous proteases.

Intraplantar administration of the PAR4 agonist peptide AYPGKF-NH_2_ (100 μg) did not significantly change withdrawal thresholds and grimacing in WT while a transient effect was seen in PAR3^-/-^ mice (**Fig. 7A-B**).

**Figure 7.**
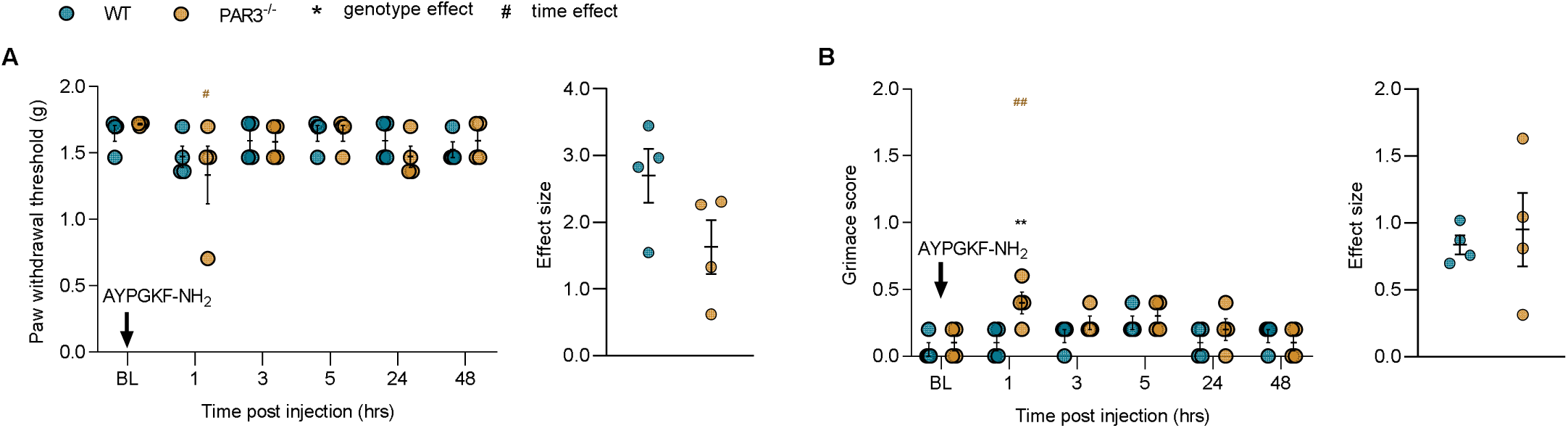
PAR4 agonist does not elicit mechanical or affective measures of pain. 100 μg of AYPGKF-NH_2_, a PAR4 agonist, was injected into the hind paw of WT and PAR3^-/-^ mice after recording baseline (BL) values. Von Frey and grimace tests were performed at 1, 3, 5, 24, and 48 hrs post injection. (A) Paw withdrawal thresholds did not change significantly in both WT and PAR3^-/-^ mice groups (n = 4/group). (B) Although facial grimacing was transiently increased with PAR3^-/-^ at 1 hr post injection, the cumulative time and genotype effects of AYPGKF-NH_2_ (100 μg, i.pl) were insignificant (n = 4/group). (Panel A and B) Data is expressed as mean ± SEM. Two-way ANOVA with Bonferroni’s multiple comparisons *p<0.05, **p<0.01. Unpaired t-test was performed for effect sizes. Stars show significant differences between treatments or genotypes. Hashtags show differences by time, from baseline.

### Hyperalgesic priming deficits in PAR3 knockout mice

Hyperalgesic priming is a two hit model where exposure to a first stimulus causes a second, normally non-noxious stimulus, to cause a long lasting pain state (Aley et al., 2000; Parada et al., 2005; Hucho and Levine, 2007). The underlying mechanisms of hyperalgesic priming involve plasticity in nociceptors (Reichling and Levine, 2009; Kandasamy and Price, 2015; Price and Inyang, 2015; Moy et al., 2017). As shown in **Fig. 6**, we observed a profound deficit in hyperalgesic priming in PAR3^-/-^ mice exposed to a PAR2 specific agonist. A potential explanation for this is a loss of PGE_2_ response in PAR3^-/-^ mice. We tested this directly by exposing mice to a high dose of PGE_2_ (10 μg). This dose of PGE_2_ caused robust mechanical hypersensitivity and grimacing in WT and PAR3^-/-^ mice (**Supplementary Fig. S5A-B**). When these mice were challenged with 100 ng PGE_2_ the animals of both genotypes displayed a response consistent with the development of hyperalgesic priming (**Supplementary Fig. S5C-D**). This shows that PAR3^-/-^ mice respond to PGE_2_ and these mice can display hyperalgesic priming, but this depends on the first hit stimulus.

To further explore which types of stimuli might show deficits in hyperalgesic priming in PAR3^-/-^ mice we assessed a number of other priming factors. The inflammatory cytokine interleukin 6 (IL-6) produced similar acute responses in both genotypes (**Fig. 8A-B**) but hyperalgesic priming was diminished as measured by mechanical hypersensitivity and grimacing in PAR3^-/-^ mice (**Fig. 8C-D**). Using carrageenan as the priming stimulus, male WT and PAR3^-/-^ mice responded similarly to the inflammagen acutely (**Supplementary Fig. S6A-B**) but the hyperalgesic priming was again reduced in the PAR3^-/-^ mice (**Supplementary Fig. S6C-D**). Similar results were obtained in female mice (**Supplementary Fig. S7A-D**). These experiments suggest that hyperalgesic priming mechanisms in response to some, but not all, priming stimuli are impaired in the absence of PAR3.

**Figure 8.**
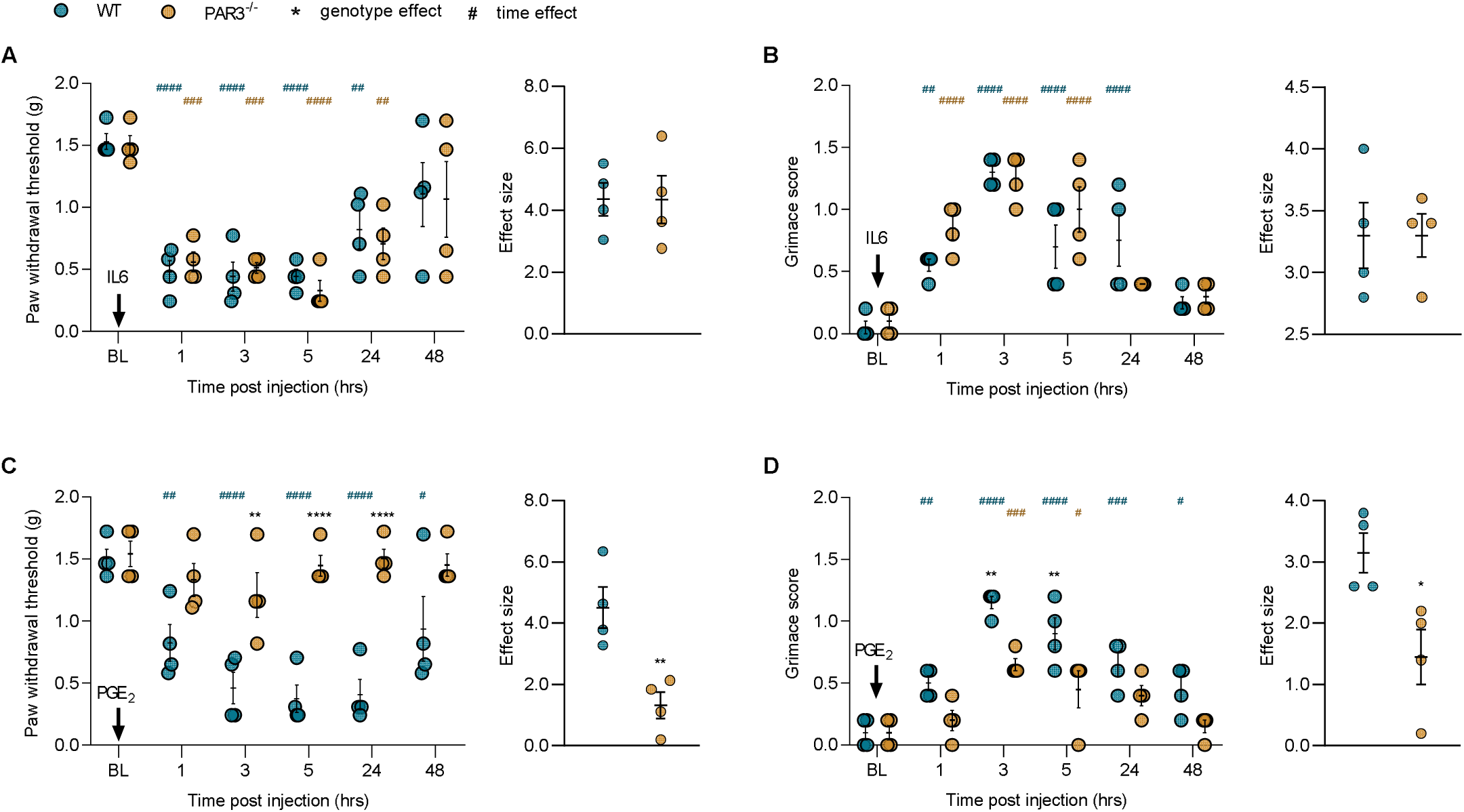
IL-6 elicits mechanical and affective pain responses but does not induce hyperalgesic priming in PAR3^-/-^ mice. (Panel A and B) WT and PAR3^-/-^ mice received an injection of IL6 (0.1 ng) into the hind paw after recording baseline (BL) values. Paw withdrawal thresholds and facial grimacing were scored at 1, 3, 5, 24, and 48 hrs post injection. (A) Both WT and PAR3^-/-^ mice demonstrated similar decreased mechanical withdrawal threshold in response to IL6 (0.1 ng, i.pl) (n = 4/group). (B) Likewise, IL6 (0.1 ng, i.pl) induced similar levels of facial grimacing in both cohorts (n = 4/group). (Panel C and D) PGE_2_ (100 ng) was administered intraplantarly in WT and PAR3^-/-^ mice, 14 d after initial stimulation with IL-6 (0.1 ng, i.pl). PAR3^-/-^ mice exhibit a deficiency in hyperalgesic priming; (C) paw withdrawal thresholds (n = 4/group) (D) and facial grimacing (n = 4/group) were not significantly affected by PGE_2_ treatment in comparison to WT mice. Data is expressed as mean ± SEM. Two-way ANOVA with Bonferroni’s multiple comparisons (Panel A-D) *p<0.05, **p<0.01, ***p<0.001, ****p<0.0001. Unpaired t-test*p<0.05, **p<0.01. Stars show significant differences between treatments or genotypes. Hashtags show differences by time, from baseline.

## DISCUSSION

Our work begins to define a role of PAR3 in nociception. PAR3 is widely distributed in mouse sensory neurons and may be crucial for inducing nociceptor hyperexcitability and mechanical hyperalgesia. We noted that PAR3 mRNA expression is detected in a majority of nociceptors regardless of the peptidergic and non-peptidergic nature of these neuronal subtypes. PAR3 expression overlaps with the presence of PAR1 or PAR2 in discrete neuronal subpopulations, likely suggesting its role in modulating PAR1- and PAR2-driven pain behaviors. We tested this hypothesis *in vivo* and observed that the involvement of PAR1 and PAR2 agonists in evoking pain stimuli are potentiated in the absence of PAR3. Critically, we have developed a novel lipid tethered peptidomimetic agonist for PAR3, C660, and demonstrated its activity and specificity both *in vitro* and *in vivo*. This tool will be useful for further understanding the role of PAR3 in pain and other areas. A remarkable phenotype of the PAR3^-/-^ mice is the loss of hyperalgesic priming in response to IL-6, carrageenan, and PAR2 agonist 2AT, suggesting that PAR3 has a role in plasticity of afferent neurons. These primary conclusions emerging from our experiments are discussed further below.

PAR3 expression has been well characterized in human and murine platelets (Ishihara et al., 1997; Ishihara et al., 1998; Schmidt et al., 1998), vascular smooth muscle cells (Bretschneider et al., 2003), endothelial cells (Schmidt et al., 1998; Cupit et al., 1999; Kalashnyk et al., 2013), and monocytes (Colognato et al., 2003), yet its presence and function in sensory neurons has not been thoroughly investigated. A previous histological study by Zhu et al. 2005 showed that, in rat DRG, PAR3 mRNA was the highest expressed of all PARs and was detected in at least 40% of neurons. In that study, they found that 80% of these PAR3 expressing cells also co-expressed CGRP (Zhu et al., 2005). Our analysis of previously published mouse single-cell RNA seq findings confirm the broad distribution patterns of PAR3 mRNA in DRG neurons and show that the mRNA is expressed in both peptidergic and non-peptidergic mouse afferents. We independently corroborated this expression pattern in mouse TG using RNAscope. Chamessian and colleagues showed that PAR3 mRNA is expressed in dorsal horn somatostatin-positive interneurons, which are known to modulate mechanical pain (Chamessian et al., 2018; Moehring et al., 2018). While we cannot confirm that we recorded from somatostatin-positive neurons in lamina II, our spinal cord slice experiments showed a clear presynaptic effect of PAR3 activation, suggesting that PAR3 expression in afferents can regulate neurotransmitter reIease onto dorsal horn neurons. Our Ca^2+^ imaging experiments on mouse DRG and TG neurons confirm that PAR3 is functionally active in at least a subset of sensory neurons. A larger population of DRG neurons expressed PAR3 mRNA than were activated by C660. This may be explained by receptor hetero-oligomerization with an unknown receptor pair or by unexplained aspects of C660 pharmacology at natively expressed PAR3. Our experiments in HEK293 cells support the conclusion that expressing PAR3 is sufficient to confer Ca^2+^ signaling in cells exposed to C660.

PAR homo- and hetero-oligomerization interactions have garnered considerable interest over the years, with several studies documenting the colocalization and transactivation of these GPCRs in different physiological settings. For instance, PAR3 associates with PAR1 in endothelial cells to potentiate the responsiveness of PAR1 to thrombin (McLaughlin et al., 2007). In other cases, PAR3 expression has been shown to counteract PAR1 signaling (Wysoczynski et al., 2010). Our results support the conclusion that PAR3 suppresses PAR1 signaling in sensory neurons. PAR1 agonists caused much larger pain responses when measured with mechanical hypersensitivity and grimacing in mice lacking PAR3. PAR2 plays a critical role in many types of persistent pain (Vergnolle et al., 2001; Lam and Schmidt, 2010; Zhao et al., 2015; Hassler et al., 2018; Jimenez-Vargas et al., 2018; Hassler et al., 2020). The specific PAR2 agonist 2AT did not show any differences in acute responses in PAR3^-/-^ mice but there was a small decrease in response to the mast cell degranulator 48/80 in these mice. This may indicate that PAR3 can regulate PAR2 activity differentially depending on the method of receptor activation. PAR3 can also complex with PAR4 in mouse platelets to facilitate the cleavage and activation of PAR4 at low thrombin concentrations (Nakanishi-Matsui et al., 2000). However, another study found that PAR3 acts as a break on PAR4 signaling in platelets (Arachiche et al., 2013), in a similar fashion to what has been described for PAR3 with PAR1 and PAR2. Nevertheless, we did not note PAR4-mediated pain behaviors in WT or PAR3^-/-^ mice suggesting that PAR4 does not play an active role in nociception from the paw. This may be different than nociception from visceral organs where PAR4 has been shown to play an important role (Bradesi, 2009; Kouzoukas et al., 2016; Wang et al., 2018). Although it is commonly thought that PAR3 cooperates with other PARs to initiate downstream signaling cascades, there is also evidence that activated PAR3 may signal autonomously to stimulate calcium mobilization and ERK1/2 phosphorylation (Bretschneider et al., 2003; Ostrowska and Reiser, 2008). Single cell sequencing data (Li et al., 2016) suggests that PAR3 is expressed in some nociceptor subtypes that do not express PAR1, -2, or -4. Therefore, we cannot exclude the possibility that PAR3 may be signaling without cooperating with other PARs in certain neuronal subpopulations. The functional role of PAR3 in those sensory neuron subtypes will need to be characterized with conditional knockout technologies.

While the oligomerization of PARs in different cell types contributes to increased receptor diversity and function, it also poses a challenge in developing agonists and antagonists that selectively act on PARs, including PAR3. To date, there are no known PAR3 antagonists and existing agonists lack potency and efficacy and have been shown to activate other PARs (Macfarlane et al., 2001; Hansen et al., 2004; Russell and McDougall, 2009; Vergnolle, 2009; Alexander et al., 2019). Using the synthetic tethered ligand approach, we have developed a more selective agonist, C660, by lipidating the peptide sequence to mimic membrane tethering that occurs with PAR endogenous ligands (Flynn et al., 2013). We show that C660 evokes Ca^2+^ responses in DRG and TG neurons and physiological responses in human PAR3-expressing cells with an EC_50_ of approximately 900 nM. Importantly, both of these responses are absent when PAR3 is not expressed. However, most DRG and TG neurons expressed PAR3 but only a subset of them showed measurable Ca^2+^ responses when C660 was applied to these cultures. We do not currently understand if PAR3 expression alone is sufficient for signaling in response to C660 although it appears to be necessary. Our *in vivo* experiments further validate the use of C660 as a pharmacological agonist of PAR3. C660 induced mechanical hypersensitivity and caused hyperalgesic priming in WT mice, but these effects were absent in PAR3 deficient mice, again supporting the specificity of this new PAR3 agonist. We anticipate that C660 will be a useful tool for further understanding the physiological role of PAR3 in different contexts and species.

Hyperalgesic priming is an animal model system used to better understand mechanisms of nociceptor plasticity that may be involved in chronic pain (Kandasamy and Price, 2015; Price and Inyang, 2015). Experimental models for hyperalgesic priming are based on the concept that the initial application of noxious stimuli may subsequently elicit prolonged pain responses to an ordinarily non-noxious stimuli (Reichling and Levine, 2009). In our experiments, we “primed” with various stimuli, including C660, 2AT, IL-6, or carrageenan allowing animals to completely recover from the initial stimulus before applying the second “hit”. PAR3 activation with C660 caused hyperalgesic priming in mice suggesting that PAR3 activation is sufficient to induce a primed state. Interestingly, in mice lacking PAR3, hyperalgesic priming failed to develop in response to most of these stimuli. This loss of hyperalgesic priming occurred in both male and female mice, at least in the carrageenan model. We acknowledge that we did not test for sex effects in most experiments, which is a shortcoming of our work. The deficit in hyperalgesic priming we observed cannot be explained by a loss of PGE_2_ sensitivity because PAR3^-/-^ responded to high dose PGE_2_ injection and even showed priming to this stimulus. Moreover, there may be differences in the role of PAR3 in priming between the DRG and TG system as we observed priming in PAR3^-/-^ mice when 2AT was applied to the dura. A caveat to this interpretation is that the second hit in the DRG (PGE_2_) and the TG (pH 7.0) were different. While we do not have a mechanistic explanation for why PAR3 appears to play a key role in nociceptor plasticity in some contexts and not in others, further investigations along these lines may reveal aspects of PAR3 signaling in nociceptors that make the receptor a drug target for chronic pain.

## MATERIALS AND METHODS

### Animals

Male, 20-30 gram mice were used in this study, ICR (Taconic, Envigo), C57BL/6J (Jackson Laboratories), and PAR3^-/-^ on a C57Bl/6J background (MMRRC, Chapel Hill, NC). The animals were housed in a climate-controlled room with a 12-hour light/dark cycle, and given food and water *ad libitum*. All experiments and procedures were performed in accordance with the guidelines recommended by the National Institute of Health, the International Association for the study of pain, and were approved by the Institutional Animal Care and Use Committee at University of Texas at Dallas or at Duke University.

### Experimental reagents

Compound 660 (TFRGAPPNSFEEF-pego3-Hdc), compound 661 (GAPPNSFEEF-pego3-Hdc), compound 662 (TFRGAP-pego3-Hdc), compound 663 (TFR-pego3-Hdc), AYPGKF-NH_2_ (PAR4 activation peptide), 2AT-LIGRL-NH_2_ (2AT), and other ligands (**Supplementary Fig. S1B**) were made using solid-phase synthesis as previously described (Boitano et al., 2011; Flynn et al., 2011). The full structure of C660 is depicted in **Supplementary Fig. S8.** Carrageenan, 48-80, thrombin, picrotoxin, strychnine and TFLLR-NH_2_ were purchased from Sigma-Aldrich, IL-6 was purchased from R&D systems; Prostaglandin E_2_ (PGE_2_) was purchased from Cayman Chemicals. Tetrodotoxin was from Tocris.

### Design of tethered PAR3 ligands

Compound 660 (TFRGAPPNSFEEF-pego3-Hdc) represents the canonical sequence of peptide TFRGAPPNSFEEF which stays tethered to the receptor activated by thrombin cleavage of the K38/T39 peptide bond in PAR3. The peptide is attached via a short trimeric pego linker to the lipid tail which resembles lipids of the cell membrane. Compound 661 (GAPPNSFEEF-pego3-Hdc), compound 662 (TFRGAP-pego3-Hdc), compound 663 (TFR-pego3-Hdc), C737 (FEEF-pego3-Hdc), and C742 (NSFEEF-pego3-Hdc) are truncated analogs of C660. Compounds C728 (Ac-pego3-Hdc) and C729 (scrambled C660 peptide PGTEFNFARESFP-pego3-Hdc) are negative controls. Compound C733 (TFRGAPPNSFEEF-amide) represents original peptide sequence without the pego linker and lipid anchor. Compound C741 (RTFRGAPPNSFEEF-pego3-Hdc) is N-terminal Arg38 extension of C660. Finally, compound C751 (TFRGAPPNSFEEF-pego3-KLIPAIYLLVFVVGV-amide) and C752 (TFRGAPPNSFEEF-pego3-KRRPAIYLLVFVVGV-amide) are analogs of C660 consisting of the active sequence TFRGAPPNSFEEF connected to the original PAR3 transmembrane peptide K^96^LIPAIYLLVFVVGV^109^ (Uniprot O00254) via a pego3 linker. We synthesized ligand with full transmembrane domain K^96^LIPAIYLLVFVVGVPANAVTLWMLF^120^, but this ligand was not tested as the lipophilic TM domain induced very poor solubility. Even compound C751 suffers from low solubility in aqueous buffers, therefore we included two arginine residues K^96^<u>RR</u>PAIYLLVFVVGV^109^ (underlined) in the transmembrane interface of C752. Solubility of C752 improved, nonetheless the compound was not active in RTCA.

### RNAscope *in situ* hybridization

For *in situ* hybridization, trigeminal ganglia (TG) from C57BL/6J male mice were dissected and post-fixed for 2 hours at 4C. The TG were cryo-sectioned to 12 μm, thaw-mounted onto Superfrost Plus (Fisher Scientific) slides, allowed to dry for 20 mins at RT, and then stored at -80C. *In situ* hybridization was performed using the RNAscope system (Advanced Cell Diagnostics). Tissue pretreatment consisted of 30 minutes of Protease IV at RT, rather than the recommended procedure for fixed frozen tissue. Following pretreatment, probe hybridization and detection with the Multiplex Fluorescence Kit v2 were performed according to the manufacturer’s protocol. Probes included Mm-F2rl2 (#489591), Mm-Trpv1-C2 (313331-C2), and Mm-Nefh-C3 (443671-C3). After detection, the tissues were counterstained with DAPI (4’,6-diamidino-2-phenylindole) and mounted with Prolong Gold (Life Technologies). Fluorescence was detected using an epifluorescence microscope (Nikon Eclipse NiE).

### Image analysis

20X images of TGs were acquired using the Nikon Eclipse NiE. Six sections of left and right TGs were imaged per mouse (n=4). Images were analysed on Olympus Cell Sens (v1.18) for the expression of *F2rl2, Trpv1* and *Nefh*. Images were first brightened and contrasted before counting the number of cells using the point tool.Total neuron counts were made for *Nefh* positive cells and all cells outlined by DAPI. F2rl2 positive neurons were then counted (including cells with less than 5 puncta) and illustrated as a percentage of the total neuronal count on a pie chart. *F2rl2* was also evaluated for co-expression with *Trpv1* in TG neurons.

### Physiological Response

96-well E-plates (ACEA Biosciences) were coated with 70μL of Poly-L-lysine (Sigma-Aldrich P4707) for 2 hours, after which Poly-L-lysine was aspirated and the plate was rinsed with sterile 18mQ water. The system set-up for the induction of PAR3 expression was adapted from a previous publication (Han et al., 2020). HEK293 cells grown at 37 °C in a 5% CO_2_ atmosphere at 70% confluence were treated with tetracycline (0.3 μg/mL or 1 μg/mL) for 48 hours to induce PAR3 expression. Cells were plated onto Poly-L-Lysine coated E-plates (ACEA Biosciences) in low serum medium (5% FBS) and monitored for Cell Index (CI) using the xCELLigence real time cell analyzer (RTCA; ACEA Biosciences) for ∼6 hours to allow for cellular attachment and stabilization. C660 was dissolved in DMEMF12 and warmed to 37 °C. For detailed protocols on running RTCA assays and interpreting CI results, please refer to (Boitano et al., 2011; Flynn et al., 2013; Boitano et al., 2015). At CI stabilization, 20 μL of 10× final concentration C660 (in DMEMF12, 37 °C) was added to each well for a final volume of 200 μL. Assays included both induced and non-induced cells treated with varying concentrations of C660 in quadruplicate. CI was measured after drug addition every minute for 4 h.

CI is dimensionless and is calculated by CI = (Z_i_ – Z_0_)/15 Ω, where Z_i_ is the impedance at an individual time point during the experiment, Z_0_ is the impedance at the start of the experiment and Ω represents ohms. This relative change in electrical impedance represents cell status, which will be affected by changes in cell morphology, adhesion or viability (Boitano et al., 2011; Flynn et al., 2013; Boitano et al., 2015).

### Analysis of in vitro physiological response

Peak changes in CI after drug addition were used to determine the dose response to C660 within one plate. These responses were normalized as percentages of the max response within each plate to determine the EC_50_ as previously described in Flynn et al. 2013 (Flynn et al., 2013). Individual traces of the CI over time represent the average of a quadruplicate from a single E-plate.

### Behavioral methods

Mice were acclimated to suspended Plexiglas chambers (9×5×5 cm high) with a wire mesh bottom (1 cm^2^). Withdrawal thresholds to probing of the hind-paws were determined before and after treatment administration. Paw withdrawal (PW) thresholds were determined by applying von Frey filaments to the plantar aspect of the hind-paws, and a response was indicated by a withdrawal of the paw. The withdrawal thresholds were determined by the Dixon up-down method (Dixon, 1980; Chaplan et al., 1994) by blinded observers.

The protocol originally developed by Mogil and colleagues for testing facial grimacing in mice was adapted for this study (Langford et al., 2010). Mice were placed individually on a tabletop in cubicles (9×5×5 cm high) with 2 walls of transparent acrylic glass and 2 side walls of removable stainless steel. Two high-resolution (1920×1080) digital video cameras (High-definition Handycam Camcorder, model HDR-CX100, Sony, San Jose, CA) were placed immediately outside both acrylic glass walls to maximize the opportunity for clear head shots. The animals were then recorded for 20 minutes and the photographs that included views of the mouse face were extracted from each recording and scored by blinded scorers. The scores were averaged at each time-point by group.

Thermal sensitivity was measured using the Hargreaves method (Hargreaves et al., 1988). Mice were placed on a heated glass floor (29°C) 20 minutes prior to each testing, and using a Hargreaves apparatus (IITC Model 390) a focused beam of high-intensity light was aimed at the plantar non-glabrous surface of the hind-paws. The intensity of the light was set to 30% of maximum with a cutoff value of 20 seconds. The latency to withdraw the hind-paw was measured to the nearest 0.01 seconds. The hind-paws were measured prior to treatment and at 1, 3, 5, 24, and 48 hours after administration.

Paw inflammation was investigated by measuring the temperature of the animal’s hind-paws. All testing was performed in a climate-controlled room with an ambient temperature of 21 ± 2°C. Animals were allowed to acclimate in the testing room for 1 hour prior to testing. Colorized infrared thermograms that captured the non-glabrous surface of the animal’s hind-paws were obtained using a FLIR T-Series Thermal Imaging Camera. The thermograms were captured prior to experimental treatment and at 1, 3, 5, 24, and 48 hours after administration. Thermogram analysis was performed using the Windows-based PC application of the FLIR system. For each thermogram image, a straight line was drawn on the plantar surface of both hind-paws and the mean temperature was recorded from the average of each pixel along the drawn line. The raw temperatures were then plotted for ipsilateral and contralateral hind-paws for each individual animal.

### Primary neuronal cultures

Male C57BL/6J or PAR3^-/-^ mice were anesthetized with isoflurane and sacrificed by decapitation. Trigeminal or dorsal root ganglia (TGs or DRGs) were dissected into Hank’s Balanced Salt Solution (HBSS), no calcium, no magnesium, on ice. Ganglia were digested in 1 mg/ml collagenase A (Roche) for 25 min at 37°C, followed by digestion in 1 mg/ml collagenase D and 30 U/ml papain (Worthington) for 20 min at 37°C. Ganglia were then triturated in 1 mg/ml trypsin inhibitor (Roche) and filtered through a 70 μm cell strainer (Corning). Cells were pelleted and resuspended in culture media, DMEM/F12 with GlutaMAX (Thermo Fisher Scientific) supplemented with 10% fetal bovine serum (FBS; SH30088.03; Hyclone) and 1% penicillin/streptomycin (Pen-Strep; 15070-063; Gibco). Cells were plated 100 uL per dish onto pre-poly-D-lysine coated dishes (P35GC-1.5-10-C; MatTek) and allowed to adhere for 2 hours before being flooded with culture media with 10 ng/ml nerve growth factor (NGF; 01-125; Millipore). The plates were kept at 37 C and 5% CO_2_ until use.

### Calcium Imaging

Ca^2+^ imaging was done using digital imaging microscopy. 48 hours after plating, the cultures were washed with HBSS and loaded in 5 μM Fura2-AM (108964-32-5; Life Technologies) in HBSS for 45 min. Fura2 fluorescence was observed on an Olympus IX70 microscope (Waltham, MA, USA) with a 40× oil immersion objective after alternating excitation between 340 and 380 nm by a 75 W Xenon lamp linked to a Delta Ram V illuminator (PTI, London, Ontario, Canada) and a gel optic line. Images were captured using a high-speed camera using Olympus software. Ca^2+^ signaling response for each individual cell in the field of view was calculated from captured images by the ratio of 340nm/380nm. A cell was considered to respond to a stimulus when there was a 10 % increase in the 340nm/380nm ratio. A minimum of one ratio per 2 seconds was calculated for all experiments.

All solutions were administered through a perfusion drip after adjusting to pH 7.4 with NaCl or N-methyl-glucamine and osmolarity to 300 ± 5 mOsm with sucrose or ddH_2_O. Normal bath solution was applied to record a stable baseline, followed by compounds at 1 uM or 100 nM in normal bath, a washout in normal bath solution, and a positive control with 50 mM KCl to identify neurons. Only cells responding to 50 mM KCl were considered for neuronal analysis. Normal bath solution consisted of NaCl (135 mM), KCl (5 mM), HEPES (10 mM), CaCl_2_ (2 μM), MgCl_2_ (1 μM) and glucose (10 μM) in ddH_2_O. KCl (50 mM) solution was made up of NaCl (90 mM), KCl (50 mM), HEPES (10 mM), CaCl_2_ (2 μM), MgCl_2_ (1 μM) and glucose (10 μM) in ddH_2_O.

### Spinal Cord slice preparation

Adult (5-7 weeks old) male C57BL/6 mice were anesthetized with urethane (1.5-2.0 g/kg, i.p.). The lumbosacral spinal cord was quickly removed and placed in ice-cold sucrose-ACSF which was saturated with 95% O_2_ and 5% CO_2_ and maintained at room temperature. After extraction and still under anesthesia, animals were sacrificed by decapitation. Transverse slices (300-400 μm) were prepared using a vibrating microslicer (VT1200s Leica). The slices were incubated at 32°C for at least 30 min in regular ACSF (NaCl 126 mM, KCl 3 mM, MgCl_2_ 1.3 mM, CaCl_2_ 2.5 mM, NaHCO_3_ 26 mM, NaH_2_PO_4_ 1.25 mM and glucose 11 mM) equilibrated with 95% O_2_ and 5% CO_2_.

### Electrophysiological recording

The slice was placed in the recording chamber and was then completely submerged and superfused at a rate of 1.5-3 ml/min with ACSF which was saturated with 95% O_2_ and 5% CO_2_ and maintained at room temperature. Lamina II neurons in lumbar segments were identified as a translucent band under a microscope (BX51WIF; Olympus) with light transmitted from below. Whole-cell voltage-clamp recordings were made from lamina II neurons by using patch-pipettes fabricated from thin-walled, fiber-filled capillaries. Patch-pipette solution used to record spontaneous excitatory postsynaptic currents (sEPSCs) contained: K-gluconate 135 mM, KCl 5 mM, CaCl_2_ 0.5 mM, MgCl_2_ 2 mM, EGTA 5 mM, HEPES 5 mM, Mg-ATP 5 mM (pH 7.3 adjusted with KOH, 300mOsm). The patch-pipettes had a resistance of 8–10 M. As previously described (Wang et al., 2020), sEPSCs recordings were made at a holding potential (V_H_) of -70 mV in the presence of 10 μM picrotoxin and 2 μM strychnine. Miniature EPSCs (mEPSCs) were recorded in the presence of 10 μM picrotoxin, 2 μM strychnine and 0.5 μM tetrodotoxin. Signals were acquired using an Axopatch 700B amplifier. The data were stored and analyzed with a personal computer using pCLAMP 10.3 software. sEPSC events were detected and analyzed using Mini Analysis Program ver. 6.0.3. Numerical data are given as the mean ± SEM. Statistical significance was determined as *P* < 0.05 using Student’s *t* test. In all cases, *n* refers to the number of the neurons studied.

### Bioinformatics

Read counts for each coding gene for 204 single cell RNA-sequencing profiles of mouse DRG sensory neurons were obtained from Gene Express Omnibus deposit (accession number GSE63576) (Li et al., 2016). t-SNE based non-linear embedding and visualization of the single cell data sets was performed using Seurat package 2.2.1 (Gribov et al., 2010; Butler et al., 2018; Hockley et al., 2019)

### Data analysis

All data are presented as means ± SEM unless otherwise noted. Unless otherwise noted, statistical evaluation was performed using two-way analysis of variance (ANOVA) with Bonferroni’s multiple comparisons. Statistical analysis was done using Graph Pad Prism Version 8.4.2 with the exception of the electrophysiology data on Figure 3 which was analysed with an earlier version (v6).

## ACKNOWLEDGEMENTS

This work was supported by NIH grants NS098826 (TJP, GD, SB and JV), NS065926 (TJP), NS072204 (GD) and training grant NS096963 (SNH).

The authors declare no conflicts of interest

