## Supplementary Information for "A role for protease activated receptor type 3 (PAR3) in nociception demonstrated through development of a novel peptide agonist"

Short Title: PAR3 contributes to nociception

Authors: Juliet Mwirigi^1*^, Moeno Kume^1*^, Shayne N Hassler^1*^, Ayesha Ahmad^1^, Pradipta R. Ray^1^, Changyu Jiang^2^, Alexander Chamessian^2^, Nakleh Mseeh^1^, Breya P Ludwig^1^, Benjamin D. Rivera^4^, Marvin T Nieman^3^, Thomas Van de Ven^2^, Ru-Rong Ji^2^, Gregory Dussor^1^, Scott Boitano^4^, Josef Vagner^5^, and Theodore J Price^1#^

^1^University of Texas at Dallas, School of Behavioral and Brain Sciences and Center for Advanced Pain Studies

^2^Duke University School of Medicine, Depts. of Anesthesiology, Pharmacology, and Cancer Biology

^3^Case Western Reserve University School of Medicine, Department of Pharmacology

^4^University of Arizona, Department of Physiology

^5^University of Arizona, Bio5 Research Institute

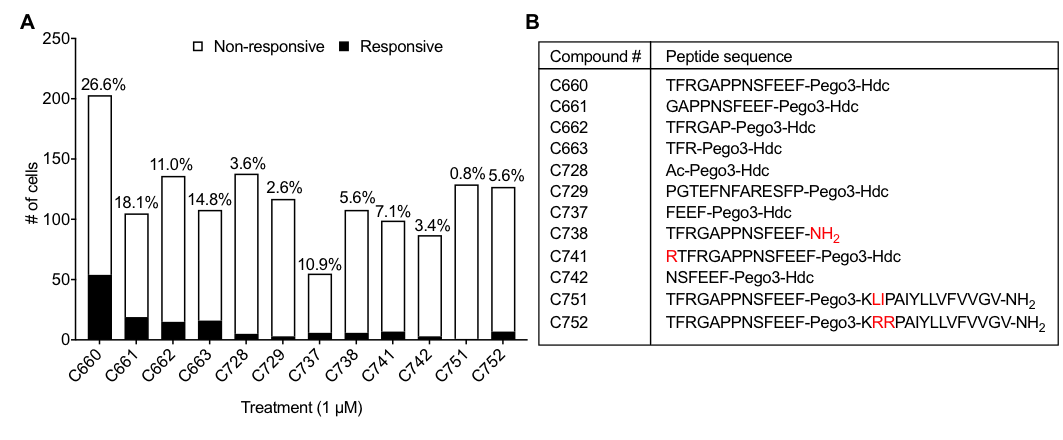

**FIG S1. C660 shows greatest activity out of compounds screened on TG cultures at 1 μM.**

Various compounds were screened for potential PAR3 activity. (A) Percent of calcium response in TG cultures at 1 μM doses of all compounds. Cells were considered responsive with a minimum 10 % increase in the 340nm/380nm ratio. C660 showed the greatest potential PAR3 activity, with a 26.6% total cell response. N = 203, 105, 136, 108, 138, 117, 55, 108, 99, 87, 129, and 127 for compounds C660-63, C728-29, C737-38, C741-42 and C751-52, respectively, where N refers to the number of cells recorded. (B) Peptide sequence of compounds screened, with red denoting the differences in structurally similar compounds. The active peptide sequence is linked to three polyethylene glycol moieties (pego3) and a hexadecyl (Hdc) lipid, except compounds C728 (plain pego3-Hdc), C738 (peptide control), and C751/ C752 (a lipid is substituted with original transmembrane peptide) .

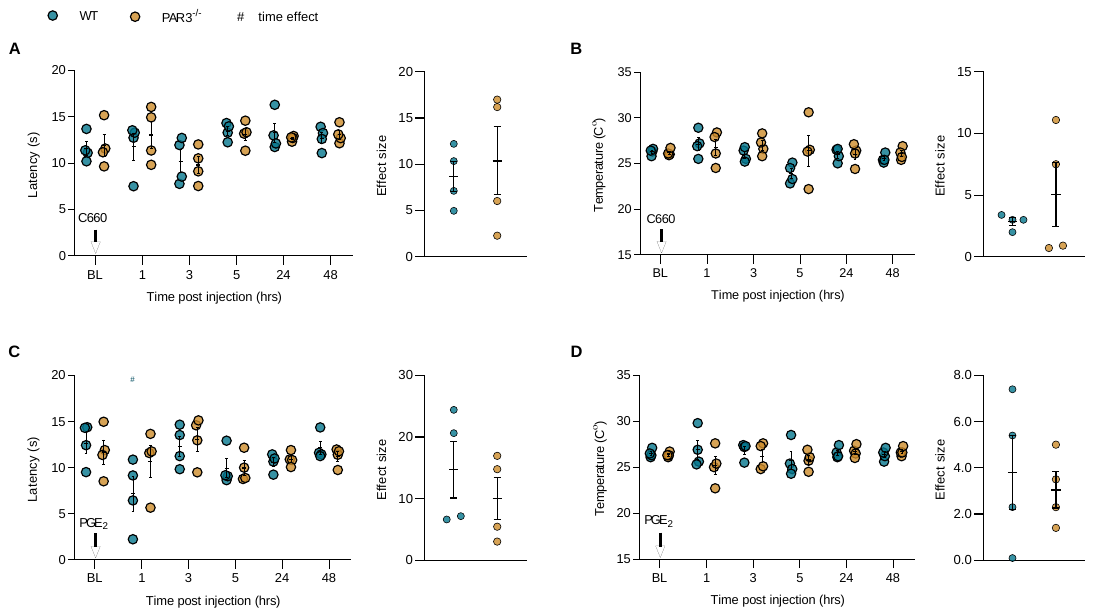

**FIG S2. Peptidomimetic agonist of PAR3 does not evoke thermal and inflammatory hyperalgesia.** (A) Mice were injected with C660 (30 pmol) into their hind paw, and thermal sensitivity was assessed via Hargreaves’ test at 1, 3, 5, 24, and 48 hours post injection. Cutoff value for paw withdrawal latency was 20 seconds. No genotype or time effect was observed in WT and PAR3^-/-^ after C660 injection (n = 4/group). (B) Likewise, assessment of paw inflammation using a FLIR thermal imaging camera showed no statistically significant genotype or time effects of C660 (30 pmol, i.pl) at 1, 3, 5, 24, and 48 hours post injection (n = 4/group). (Panel C and D) PGE_2_ (100 ng) was injected intraplantarly in WT and PAR3^-/-^ mice 14 days after initial stimulation with C660 (30 pmol). Withdrawal latency and paw inflammation were assessed 1, 3, 5, 24, and 48 hrs post PGE_2_ injection via Hargreaves’ test and FLIR analysis, respectively. (C) A transient decrease in paw withdrawal latency was observed with the WT group at the 1 hr timepoint only. There were no significant genotypic differences between the WT and PAR3^-/-^ mice in response to PGE_2_ in the hyperalgesic priming paradigm (n = 4/group). (D) Similarly, hind paw temperatures recorded post PGE_2_ (100 ng, i.pl) injection show no differences in genotype at the indicated timepoints (n = 4/group). (Panel A-D) Baseline (BL) measures were recorded beforehand. Data is expressed as mean ± SEM. Two-way ANOVA with Bonferroni’s multiple comparisons *p<0.05. Unpaired t-test was used to statistically analyze the effect sizes. Hashtags show differences by time, from baseline.

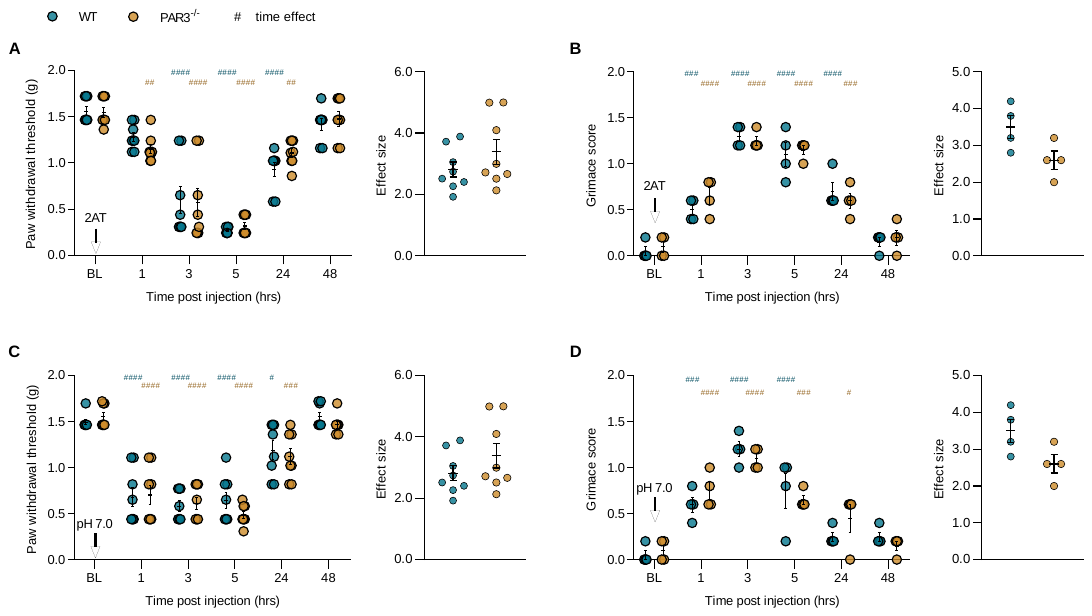

**FIG S3. Supradural administration of 2AT evokes hyperalgesia and hyperalgesic priming in both WT and PAR3^-/-^ mice.** (Panel A and B) PAR2 agonist, 2AT (30 pmol), was applied to the dural meninges of WT and PAR3^-/-^ mice. Mechanical and affective measures of pain were subsequently scored at 1, 3, 5, 24, and 48 hours post injection. 2AT (30 pmol) significantly induced (A) mechanical hypersensitivity and (B) grimacing in both WT and PAR3^-/-^ mice, with effects lasting up to 24 hrs post injection (n = 4/group) and no genotypic differences. (Panel C and D) Paw withdrawal thresholds and facial grimacing were scored after supradural application of pH 7.0 vehicle (5 µl) in WT and PAR3^-/-^ mice 14 days post initial stimulation with 2AT. pH 7.0 demonstrate hyperalgesic priming in response to 2AT as demonstrated by a significant decrease in (C) paw withdrawal thresholds up to 24 hours for both groups and an increase in (D) facial grimacing for the duration of 5 hrs with WT and 24 hrs with PAR3^-/-^ mice. (Panel A-D) Data is expressed as mean ± SEM. Two-way ANOVA with Bonferroni’s multiple comparisons *p<0.05, **p<0.01, ***p<0.001, ****p<0.0001. Unpaired t-test for effect sizes yielded non-significant results. Hashtags show differences by time, from baseline.

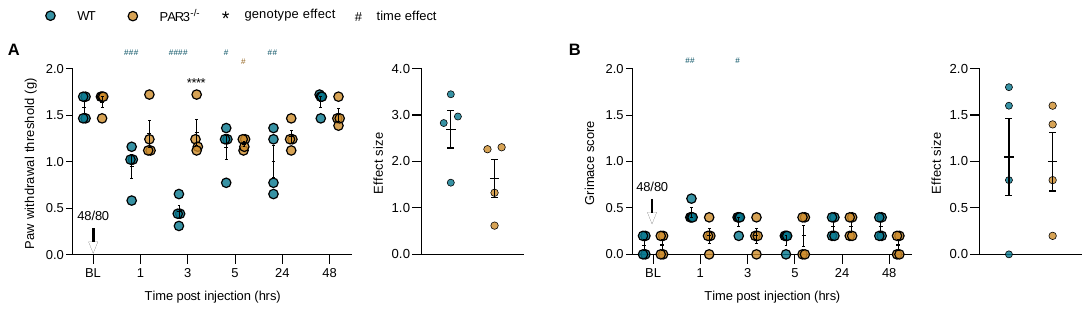

**FIG S4. Mast cell degranulator 48/80 does produce hyperalgesia in PAR3^-/-^ mice.** 48/80 (6.5 nmol) was administered interplantarly to WT and PAR3^-/-^ mice after assessing baseline measures (BL) for paw withdrawal and grimacing. (A) 48/80 significantly reduced paw withdrawal thresholds in WT at 1-, 3-, 5-, and 24 hours post injection. Genotypic differences between WT and PAR3^-/-^ was observed, particularly at the 3hr time point, but the overall effect size was not significant (n = 4/group). (B) 48/80 (6.5 nmol) induced nocifensive facial expressions in WT mice only at 1- and 3 hrs post injection (n = 4/group). (A and B) Data is expressed as mean ± SEM. Two-way ANOVA with Bonferroni’s multiple comparisons *p<0.05, **p<0.01, ***p<0.001, ****p<0.0001. Unpaired t-test for effect sizes yielded nonsignificant results. Stars show significant differences between treatments or genotypes. Hashtags show differences by time, from baseline.

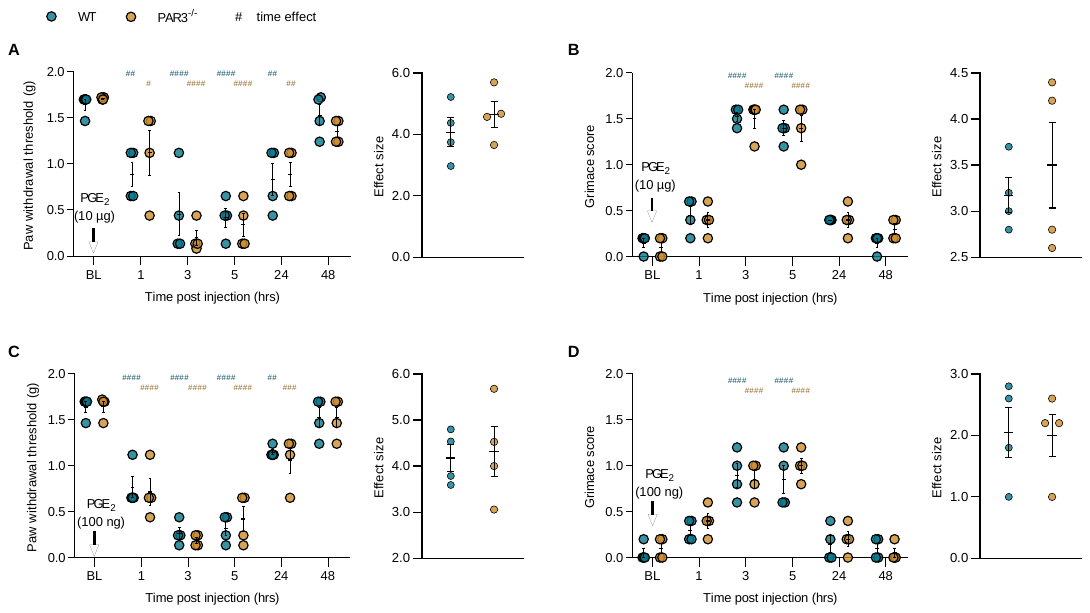

**FIG S5. PGE_2_-evoked pain behaviors.** (Panel A and B) WT and PAR3^-/-^ mice received an intraplantar injection of PGE_2_ (10 µg) after assessing baseline (BL) values. PGE_2_ (10 µg, i.pl) elicits robust mechanical and affective pain behaviors in both WT and PAR3^-/-^; (B) mechanical hypersensitivity (n = 4/group) and (C) facial grimacing (n = 4/group) was observed at 1 hr post injection, with effects lasting up to 24 hrs. (Panel C and D) Application of a normally non-noxious dose of PGE_2_ (100 ng, i.pl), 14 days after priming stimulation with PGE_2_ (10 µg, i.pl) evoked (C) mechanical hypersensitivity (n = 4/group) and (D) facial grimacing (n = 4/group) in both WT and PAR3^-/-^. Data is expressed as mean ± SEM. Two-way ANOVA with Bonferroni’s multiple comparisons (Panel A-D) *p<0.05, **p<0.01, ***p<0.001, ****p<0.0001. Hashtags show differences by time, from baseline.

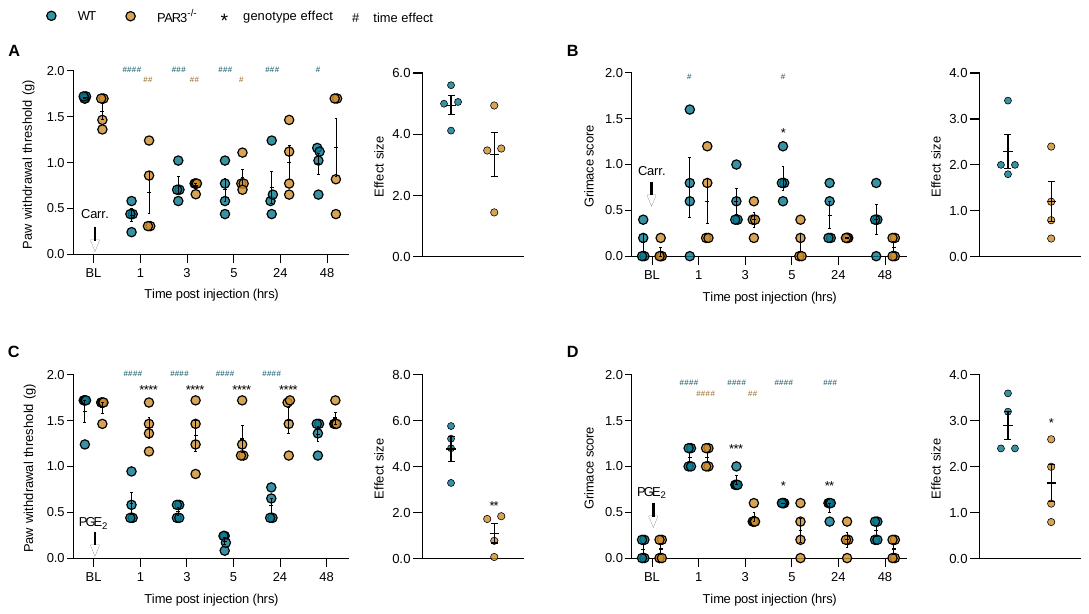

**FIG S6. Absence of carrageenan-evoked hyperalgesic priming in male mice.** Carrageenan (2%, 25 µL) was injected into the hind paw of male WT and PAR3^-/-^ mice. Paw withdrawal thresholds and facial grimacing were subsequently scored at 1, 3, 5, 24 and 48 hours post injection, and after obtaining baseline (BL) values (A) Carrageenan triggered mechanical hypersensitivity in WT and PAR3^-/-^ mice, and the cumulative genotype differences were not significant (n = 4/group). (B) Facial grimacing in response to carrageenan was observed with the WT cohort, 1 and 3 hrs post injection (n = 4/group). (C, D) The hyperalgesic priming effect of carrageenan (2%, 25 µL, i.pl) was tested on male WT and PAR3^-/-^ by injecting PGE_2_ (100ng, i.pl), 14 days later. Hyperalgesic priming present in the WT but not PAR3^-/-^ mice (n = 4/group). Data is expressed as mean ± SEM. Two-way ANOVA with Bonferroni’s multiple comparisons (Panel A-D) *p<0.05, **p<0.01, ***p<0.001, ****p<0.0001. Effect sizes were analyzed using the unpaired t-test *p<0.05, **p<0.01. Stars show significant differences between treatments or genotypes. Hashtags show differences by time, from baseline.

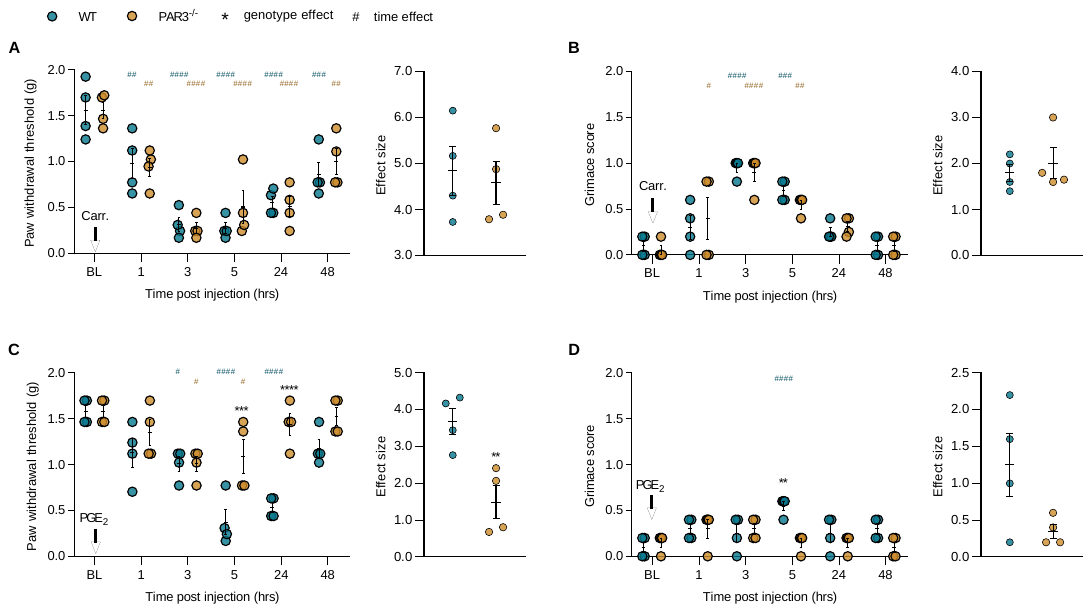

**FIG S7. Absence of carrageenan-evoked hyperalgesic priming in female mice.** Carrageenan (2%, 25 µL) was injected into the hind paw of female WT and PAR3^-/-^ mice after assessing baseline (BL) measures. (A) A robust and prolonged hyperalgesic response to carrageenan (2%, 25 µL, i.pl) was present in both WT and PAR3^-/-^ groups with no significant genotype differences (n = 4/group). (B) Carrageenan effectuated facial grimacing in WT females at 3 and 5 hrs post injection and in PAR3^-/-^ females at 1, 3, and 5 hrs post injection (n = 4/group) (C, D). Fourteen days later PGE_2_ (100 ng) was injected interplantarly, after which paw withdrawal thresholds and grimace scores were assessed at 1, 3, 5, 24 and 48 hours post injection. (C) Loss of PAR3 inhibited carrageenan-induced hyperalgesic priming (n = 4/group). (D) Cumulatively, facial grimacing was not significantly affected in WT and PAR3^-/-^(n = 4/group). Data is expressed as mean ± SEM. Two-way ANOVA with Bonferroni’s multiple comparisons (Panel A-D) *p<0.05, **p<0.01, ***p<0.001, ****p<0.0001. Effect sizes were analyzed using the unpaired t-test, **p<0.01. Stars show significant differences between treatments or genotypes. Hashtags show differences by time, from baseline.

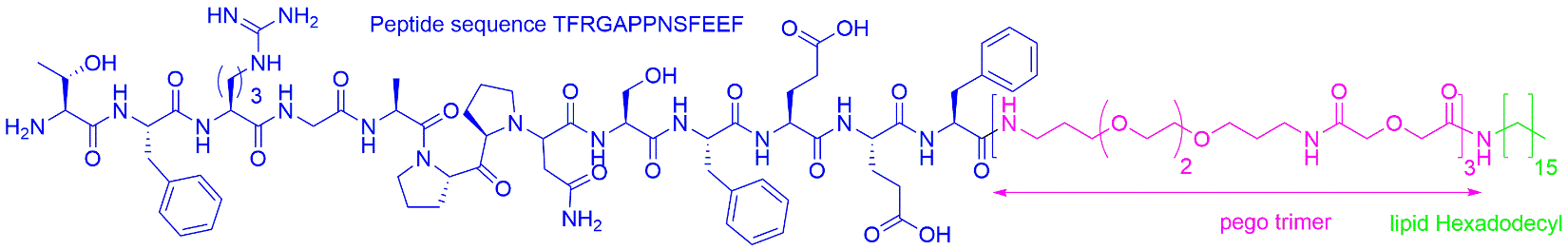

**FIG S8. Full structure of C660 compound.**

Canonical peptide structure of tethered peptide Thr39-Phe51 of human PAR3 receptor is colored in blue. Short polyethyelene glycol linker pego3 is pink and lipid moiety is green

**Supplementary Table 1** - Two-way ANOVA with Bonferroni’s Multiple Comparisons, paired and unpaired t-tests

| Fig | | **Agonist dose-response curve** | | | | | | | | | | | | | | | | |
| --- | --- | --- | --- | --- | --- | --- | --- | --- | --- | --- | --- | --- | --- | --- | --- | --- | --- | --- |
| **2E** | | C660 EC_50_(95% CI) | | | | | | | | 8.812e-007 (2.816e-008 -2.757e-005) | | | | | | | | |
| **Paired t-test** | | | | | | | | | | | | | | | | | | |
| Fig | | (t,DF), p | | | | | | | | Fig | | (t,DF), p | | | | | | |
| **3B** | | 5.361 | | 4 | | | | 0.0058 | | **3C** | | 1.196 | | 4 | | | 0.2977 | |
| **3E** | | 4.631 | | 3 | | | | 0.0190 | | **3F** | | 0.6473 | | 3 | | | 0.5636 | |
| **Two-way ANOVA Bonferroni’s Multiple Comparisons** | | | | | | | | | | | | | | | | | | |
| Fig | Comparison | | | | (t,DF), p | | | | | Fig | Comparison | | | | (t,DF), p | | | |
|  | WT - PAR3^-/-^ | | | | | | | | |  | WT - PAR3^-/-^ | | | | | | | |
|  |  | | | | t | DF | | | p |  |  | | | | t | DF | | p |
| **4A** | BL | | | | 0.4680 | 36.00 | | | >0.9999 | **6B** | BL | | | | 0.5000 | 36.00 | | >0.9999 |
|  | Hour 1 | | | | 3.977 | 36.00 | | | 0.0019 |  | Hour 1 | | | | 0.000 | 36.00 | | >0.9999 |
|  | Hour 3 | | | | 7.299 | 36.00 | | | <0.0001 |  | Hour 3 | | | | 2.000 | 36.00 | | 0.3185 |
|  | Hour 5 | | | | 7.716 | 36.00 | | | <0.0001 |  | Hour 5 | | | | 0.5000 | 36.00 | | >0.9999 |
|  | Hour 24 | | | | 8.945 | 36.00 | | | <0.0001 |  | Hour 24 | | | | 0.5000 | 36.00 | | >0.9999 |
|  | Hour 48 | | | | 2.494 | 36.00 | | | 0.1041 |  | Hour 48 | | | | 0.5000 | 36.00 | | >0.9999 |
| **4B** | BL | | | | 0.6367 | 30.00 | | | >0.9999 | **6C** | BL | | | | 0.2423 | 36.00 | | >0.9999 |
|  | Hour 1 | | | | 4.457 | 30.00 | | | 0.0005 |  | Hour 1 | | | | 6.178 | 36.00 | | <0.0001 |
|  | Hour 3 | | | | 1.273 | 30.00 | | | >0.9999 |  | Hour 3 | | | | 6.583 | 36.00 | | <0.0001 |
|  | Hour 5 | | | | 0.6367 | 30.00 | | | >0.9999 |  | Hour 5 | | | | 8.641 | 36.00 | | <0.0001 |
|  | Hour 24 | | | | 0.000 | 30.00 | | | >0.9999 |  | Hour 24 | | | | 8.501 | 36.00 | | <0.0001 |
|  |  | | | |  |  | | |  |  | Hour 48 | | | | 1.995 | 36.00 | | 0.3220 |
| **4C** | BL | | | | 0.4069 | 36.00 | | | >0.9999 | **6D** | BL | | | | 0.000 | 36.00 | | >0.9999 |
|  | Hour 1 | | | | 5.337 | 36.00 | | | <0.0001 |  | Hour 1 | | | | 2.058 | 36.00 | | 0.2813 |
|  | Hour 3 | | | | 3.940 | 36.00 | | | 0.0022 |  | Hour 3 | | | | 2.058 | 36.00 | | 0.2813 |
|  | Hour 5 | | | | 3.846 | 36.00 | | | 0.0028 |  | Hour 5 | | | | 1.543 | 36.00 | | 0.7888 |
|  | Hour 24 | | | | 0.9418 | 36.00 | | | >0.9999 |  | Hour 24 | | | | 1.543 | 36.00 | | 0.7888 |
|  | Hour 48 | | | | 2.180 | 36.00 | | | 0.2151 |  | Hour 48 | | | | 2.058 | 36.00 | | 0.2813 |
| **4D** | BL | | | | 0.5941 | 36.00 | | | >0.9999 | **7A** | BL | | | | 0.5525 | 36.00 | | >0.9999 |
|  | Hour 1 | | | | 2.970 | 36.00 | | | 0.0316 |  | Hour 1 | | | | 1.096 | 36.00 | | >0.9999 |
|  | Hour 3 | | | | 8.317 | 36.00 | | | <0.0001 |  | Hour 3 | | | | 0.08929 | 36.00 | | >0.9999 |
|  | Hour 5 | | | | 2.970 | 36.00 | | | 0.0316 |  | Hour 5 | | | | 0.000 | 36.00 | | >0.9999 |
|  | Hour 24 | | | | 0.5941 | 36.00 | | | >0.9999 |  | Hour 24 | | | | 0.9696 | 36.00 | | >0.9999 |
|  | Hour 48 | | | | 0.5941 | 36.00 | | | >0.9999 |  | Hour 48 | | | | 0.5525 | 36.00 | | >0.9999 |
| **5A** | BL | | | | 0.3039 | 36.00 | | | >0.9999 | **7B** | BL | | | | 0.5941 | 36.00 | | >0.9999 |
|  | Hour 1 | | | | 5.392 | 36.00 | | | <0.0001 |  | Hour 1 | | | | 3.565 | 36.00 | | 0.0063 |
|  | Hour 3 | | | | 3.757 | 36.00 | | | 0.0037 |  | Hour 3 | | | | 1.188 | 36.00 | | >0.9999 |
|  | Hour 5 | | | | 2.854 | 36.00 | | | 0.0427 |  | Hour 5 | | | | 0.5941 | 36.00 | | >0.9999 |
|  | Hour 24 | | | | 2.471 | 36.00 | | | 0.1099 |  | Hour 24 | | | | 1.188 | 36.00 | | >0.9999 |
|  | Hour 48 | | | | 1.750 | 36.00 | | | 0.5322 |  | Hour 48 | | | | 0.5941 | 36.00 | | >0.9999 |
| **5B** | BL | | | | 0.4271 | 36.00 | | | >0.9999 | **8A** | BL | | | | 0.1283 | 36.00 | | >0.9999 |
|  | Hour 1 | | | | 2.135 | 36.00 | | | 0.2376 |  | Hour 1 | | | | 0.3858 | 36.00 | | >0.9999 |
|  | Hour 3 | | | | 8.969 | 36.00 | | | <0.0001 |  | Hour 3 | | | | 0.3438 | 36.00 | | >0.9999 |
|  | Hour 5 | | | | 4.271 | 36.00 | | | 0.0008 |  | Hour 5 | | | | 0.5645 | 36.00 | | >0.9999 |
|  | Hour 24 | | | | 1.281 | 36.00 | | | >0.9999 |  | Hour 24 | | | | 0.5623 | 36.00 | | >0.9999 |
|  | Hour 48 | | | | 2.456 | 36.00 | | | 0.1141 |  | Hour 48 | | | | 0.1987 | 36.00 | | >0.9999 |
| **5C** | BL | | | | 0.5069 | 36.00 | | | >0.9999 | **8B** | BL | | | | 0.3254 | 36.00 | | >0.9999 |
|  | Hour 1 | | | | 7.531 | 36.00 | | | <0.0001 |  | Hour 1 | | | | 1.952 | 36.00 | | 0.3522 |
|  | Hour 3 | | | | 5.015 | 36.00 | | | <0.0001 |  | Hour 3 | | | | 0.3254 | 36.00 | | >0.9999 |
|  | Hour 5 | | | | 5.767 | 36.00 | | | <0.0001 |  | Hour 5 | | | | 1.952 | 36.00 | | 0.3522 |
|  | Hour 24 | | | | 6.838 | 36.00 | | | <0.0001 |  | Hour 24 | | | | 2.278 | 36.00 | | 0.1727 |
|  | Hour 48 | | | | 1.712 | 36.00 | | | 0.5729 |  | Hour 48 | | | | 0.3254 | 36.00 | | >0.9999 |
| **5D** | BL | | | | 2.081e-015 | 36.00 | | | >0.9999 | **8C** | BL | | | | 0.1960 | 36.00 | | >0.9999 |
|  | Hour 1 | | | | 0.4685 | 36.00 | | | >0.9999 |  | Hour 1 | | | | 2.652 | 36.00 | | 0.0709 |
|  | Hour 3 | | | | 7.028 | 36.00 | | | <0.0001 |  | Hour 3 | | | | 3.907 | 36.00 | | 0.0024 |
|  | Hour 5 | | | | 6.091 | 36.00 | | | <0.0001 |  | Hour 5 | | | | 5.584 | 36.00 | | <0.0001 |
|  | Hour 24 | | | | 4.685 | 36.00 | | | 0.0002 |  | Hour 24 | | | | 5.712 | 36.00 | | <0.0001 |
|  | Hour 48 | | | | 2.343 | 36.00 | | | 0.1488 |  | Hour 48 | | | | 2.675 | 36.00 | | 0.0670 |
| **6A** | BL | | | | 0.1910 | 36.00 | | | >0.9999 | **8D** | BL | | | | 1.830e-015 | 36.00 | | >0.9999 |
|  | Hour 1 | | | | 2.106 | 36.00 | | | 0.2535 |  | Hour 1 | | | | 2.472 | 36.00 | | 0.1097 |
|  | Hour 3 | | | | 0.2712 | 36.00 | | | >0.9999 |  | Hour 3 | | | | 4.121 | 36.00 | | 0.0013 |
|  | Hour 5 | | | | 0.07329 | 36.00 | | | >0.9999 |  | Hour 5 | | | | 3.709 | 36.00 | | 0.0042 |
|  | Hour 24 | | | | 1.058 | 36.00 | | | >0.9999 |  | Hour 24 | | | | 2.060 | 36.00 | | 0.2799 |
|  | Hour 48 | | | | 0.6889 | 36.00 | | | >0.9999 |  | Hour 48 | | | | 2.472 | 36.00 | | 0.1097 |
| **Unpaired t-test** | | | | | | | | | | | | | | | | | | |
| Fig | (t,DF), p | | | | | | | | | Fig | (t,DF), p | | | | | | | |
| **4A** | 8.098 | | 6 | | | | 0.0002 | | | **6B** | 0.2756 | | 6 | | | 0.7921 | | |
| **4B** | 0.4714 | | 6 | | | | 0.6540 | | | **6C** | 7.347 | | 6 | | | 0.0003 | | |
| **4C** | 2.224 | | 6 | | | | 0.0678 | | | **6D** | 1.782 | | 6 | | | 0.1250 | | |
| **4D** | 2.733 | | 6 | | | | 0.0340 | | | **7A** | 0.3999 | | 6 | | | 0.7031 | | |
| **5A** | 4.292 | | 6 | | | | 0.0051 | | | **7B** | 0.2000 | | 6 | | | 0.8481 | | |
| **5B** | 2.997 | | 6 | | | | 0.0241 | | | **8A** | 0.009927 | | 6 | | | 0.9924 | | |
| **5C** | 10.36 | | 6 | | | | <0.0001 | | | **8B** | 0.000 | | 6 | | | >0.9999 | | |
| **5D** | 4.317 | | 6 | | | | 0.0050 | | | **8C** | 3.996 | | 6 | | | 0.0072 | | |
| **6A** | 0.4635 | | 6 | | | | 0.6594 | | | **8D** | 3.078 | | 6 | | | 0.0217 | | |

**Supplementary Table 2** – two-way ANOVA with Bonferroni’s Multiple Comparisons

| Fig | Comparison | (t,DF), p | | | | Comparison | (t,DF), p | | |
| --- | --- | --- | --- | --- | --- | --- | --- | --- | --- |
|  | WT | | | | | PAR3^-/-^ | | | |
|  |  | t | DF | p | |  | t | DF | p |
| **4A** | BL vs. Hour 1 | 7.560 | 30.00 | <0.00011 | | BL vs. Hour 1 | 3.871 | 30.00 | 0.0027 |
|  | BL vs. Hour 3 | 12.33 | 30.00 | <0.0001 | | BL vs. Hour 3 | 5.150 | 30.00 | <0.0001 |
|  | BL vs. Hour 5 | 11.04 | 30.00 | <0.0001 | | BL vs. Hour 5 | 3.417 | 30.00 | 0.0092 |
|  | BL vs. Hour 24 | 12.10 | 30.00 | <0.0001 | | BL vs. Hour 24 | 3.189 | 30.00 | 0.0166 |
|  | BL vs. Hour 48 | 4.131 | 30.00 | 0.0013 | | BL vs. Hour 48 | 2.000 | 30.00 | 0.2732 |
| **4B** | BL vs. Hour 1 | 3.835 | 24.00 | 0.0032 | | BL vs. Hour 1 | 0.7670 | 24.00 | >0.9999 |
|  | BL vs. Hour 3 | 1.534 | 24.00 | 0.5525 | | BL vs. Hour 3 | 0.7670 | 24.00 | >0.9999 |
|  | BL vs. Hour 5 | 0.000 | 24.00 | >0.9999 | | BL vs. Hour 5 | 0.000 | 24.00 | >0.9999 |
|  | BL vs. Hour 24 | 1.534 | 24.00 | 0.5525 | | BL vs. Hour 24 | 0.7670 | 24.00 | >0.9999 |
| **4C** | BL vs. Hour 1 | 8.761 | 30.00 | <0.0001 | | BL vs. Hour 1 | 2.958 | 30.00 | 0.0300 |
|  | BL vs. Hour 3 | 8.054 | 30.00 | <0.0001 | | BL vs. Hour 3 | 3.663 | 30.00 | 0.0048 |
|  | BL vs. Hour 5 | 6.500 | 30.00 | <0.0001 | | BL vs. Hour 5 | 2.203 | 30.00 | 0.1771 |
|  | BL vs. Hour 24 | 1.363 | 30.00 | 0.9158 | | BL vs. Hour 24 | 0.000 | 30.00 | >0.9999 |
|  | BL vs. Hour 48 | 0.9959 | 30.00 | >0.9999 | | BL vs. Hour 48 | 0.7959 | 30.00 | >0.9999 |
| **4D** | BL vs. Hour 1 | 5.265 | 30.00 | | <0.0001 | BL vs. Hour 1 | 1.755 | 30.00 | 0.4474 |
|  | BL vs. Hour 3 | 11.11 | 30.00 | | <0.0001 | BL vs. Hour 3 | 2.340 | 30.00 | 0.1307 |
|  | BL vs. Hour 5 | 4.095 | 30.00 | | 0.0015 | BL vs. Hour 5 | 0.5850 | 30.00 | >0.9999 |
|  | BL vs. Hour 24 | 1.170 | 30.00 | | >0.9999 | BL vs. Hour 24 | 6.495e-016 | 30.00 | >0.9999 |
|  | BL vs. Hour 48 | 6.495e-016 | 30.00 | | >0.9999 | BL vs. Hour 48 | 6.495e-016 | 30.00 | >0.9999 |
| **5A** | BL vs. Hour 1 | 2.014 | 30.00 | 0.2653 | | BL vs. Hour 1 | 8.697 | 30.00 | <0.0001 |
|  | BL vs. Hour 3 | 3.234 | 30.00 | 0.0148 | | BL vs. Hour 3 | 7.998 | 30.00 | <0.0001 |
|  | BL vs. Hour 5 | 4.432 | 30.00 | 0.0006 | | BL vs. Hour 5 | 8.138 | 30.00 | <0.0001 |
|  | BL vs. Hour 24 | 3.548 | 30.00 | 0.0065 | | BL vs. Hour 24 | 6.804 | 30.00 | <0.0001 |
|  | BL vs. Hour 48 | 3.804 | 30.00 | 0.0033 | | BL vs. Hour 48 | 6.213 | 30.00 | <0.0001 |
| **5B** | BL vs. Hour 1 | 2.658 | 30.00 | 0.0623 | | BL vs. Hour 1 | 0.000 | 30.00 | >0.9999 |
|  | BL vs. Hour 3 | 1.329 | 30.00 | 0.9690 | | BL vs. Hour 3 | 10.19 | 30.00 | <0.0001 |
|  | BL vs. Hour 5 | 2.658 | 30.00 | 0.0623 | | BL vs. Hour 5 | 6.646 | 30.00 | <0.0001 |
|  | BL vs. Hour 24 | 2.215 | 30.00 | 0.1723 | | BL vs. Hour 24 | 3.102 | 30.00 | 0.0208 |
|  | BL vs. Hour 48 | 0.5538 | 30.00 | >0.9999 | | BL vs. Hour 48 | 2.658 | 30.00 | 0.0623 |
| **5C** | BL vs. Hour 1 | 5.306 | 30.00 | <0.0001 | | BL vs. Hour 1 | 12.40 | 30.00 | <0.0001 |
|  | BL vs. Hour 3 | 8.007 | 30.00 | <0.0001 | | BL vs. Hour 3 | 12.56 | 30.00 | <0.0001 |
|  | BL vs. Hour 5 | 5.596 | 30.00 | <0.0001 | | BL vs. Hour 5 | 10.91 | 30.00 | <0.0001 |
|  | BL vs. Hour 24 | 4.089 | 30.00 | 0.0015 | | BL vs. Hour 24 | 10.48 | 30.00 | <0.0001 |
|  | BL vs. Hour 48 | 1.927 | 30.00 | 0.3173 | | BL vs. Hour 48 | 3.145 | 30.00 | 0.0187 |
| **5D** | BL vs. Hour 1 | 3.540 | 30.00 | 0.0066 | | BL vs. Hour 1 | 3.034 | 30.00 | 0.0247 |
|  | BL vs. Hour 3 | 3.540 | 30.00 | 0.0066 | | BL vs. Hour 3 | 11.12 | 30.00 | <0.0001 |
|  | BL vs. Hour 5 | 2.528 | 30.00 | 0.0848 | | BL vs. Hour 5 | 9.102 | 30.00 | <0.0001 |
|  | BL vs. Hour 24 | 1.517 | 30.00 | 0.6987 | | BL vs. Hour 24 | 6.573 | 30.00 | <0.0001 |
|  | BL vs. Hour 48 | 1.011 | 30.00 | >0.9999 | | BL vs. Hour 48 | 3.540 | 30.00 | 0.0066 |
| **6A** | BL vs. Hour 1 | 6.570 | 30.00 | <0.0001 | | BL vs. Hour 1 | 4.674 | 30.00 | 0.0003 |
|  | BL vs. Hour 3 | 6.162 | 30.00 | <0.0001 | | BL vs. Hour 3 | 6.083 | 30.00 | <0.0001 |
|  | BL vs. Hour 5 | 6.456 | 30.00 | <0.0001 | | BL vs. Hour 5 | 6.573 | 30.00 | <0.0001 |
|  | BL vs. Hour 24 | 3.378 | 30.00 | 0.0102 | | BL vs. Hour 24 | 2.520 | 30.00 | 0.0865 |
|  | BL vs. Hour 48 | 0.1787 | 30.00 | >0.9999 | | BL vs. Hour 48 | 1.050 | 30.00 | >0.9999 |
| **6B** | BL vs. Hour 1 | 3.946 | 30.00 | 0.0022 | | BL vs. Hour 1 | 3.452 | 30.00 | 0.0084 |
|  | BL vs. Hour 3 | 3.946 | 30.00 | 0.0022 | | BL vs. Hour 3 | 5.425 | 30.00 | <0.0001 |
|  | BL vs. Hour 5 | 8.384 | 30.00 | <0.0001 | | BL vs. Hour 5 | 7.398 | 30.00 | <0.0001 |
|  | BL vs. Hour 24 | 1.973 | 30.00 | 0.2890 | | BL vs. Hour 24 | 1.973 | 30.00 | 0.2890 |
|  | BL vs. Hour 48 | 0.9864 | 30.00 | >0.9999 | | BL vs. Hour 48 | 0.000 | 30.00 | >0.9999 |
| **6C** | BL vs. Hour 1 | 5.880 | 30.00 | <0.0001 | | BL vs. Hour 1 | 0.04180 | 30.00 | >0.9999 |
|  | BL vs. Hour 3 | 7.174 | 30.00 | <0.0001 | | BL vs. Hour 3 | 0.9378 | 30.00 | >0.9999 |
|  | BL vs. Hour 5 | 9.171 | 30.00 | <0.0001 | | BL vs. Hour 5 | 0.9091 | 30.00 | >0.9999 |
|  | BL vs. Hour 24 | 7.635 | 30.00 | <0.0001 | | BL vs. Hour 24 | 0.4885 | 30.00 | >0.9999 |
|  | BL vs. Hour 48 | 2.577 | 30.00 | 0.0757 | | BL vs. Hour 48 | 0.8529 | 30.00 | >0.9999 |
| **6D** | BL vs. Hour 1 | 4.243 | 30.00 | 0.0010 | | BL vs. Hour 1 | 2.121 | 30.00 | 0.2113 |
|  | BL vs. Hour 3 | 7.955 | 30.00 | <0.0001 | | BL vs. Hour 3 | 5.834 | 30.00 | <0.0001 |
|  | BL vs. Hour 5 | 5.303 | 30.00 | <0.0001 | | BL vs. Hour 5 | 6.894 | 30.00 | <0.0001 |
|  | BL vs. Hour 24 | 4.773 | 30.00 | 0.0002 | | BL vs. Hour 24 | 3.182 | 30.00 | 0.0170 |
|  | BL vs. Hour 48 | 2.121 | 30.00 | 0.2113 | | BL vs. Hour 48 | 5.888e-016 | 30.00 | >0.9999 |
| **7A** | BL vs. Hour 1 | 1.349 | 30.00 | 0.9366 | | BL vs. Hour 1 | 2.952 | 30.00 | 0.0304 |
|  | BL vs. Hour 3 | 0.4068 | 30.00 | >0.9999 | | BL vs. Hour 3 | 1.031 | 30.00 | >0.9999 |
|  | BL vs. Hour 5 | 9.675e-006 | 30.00 | >0.9999 | | BL vs. Hour 5 | 0.5370 | 30.00 | >0.9999 |
|  | BL vs. Hour 24 | 0.4068 | 30.00 | >0.9999 | | BL vs. Hour 24 | 1.886 | 30.00 | 0.3448 |
|  | BL vs. Hour 48 | 0.9439 | 30.00 | >0.9999 | | BL vs. Hour 48 | 0.9439 | 30.00 | >0.9999 |
| **7B** | BL vs. Hour 1 | 0.5941 | 30.00 | >0.9999 | | BL vs. Hour 1 | 3.565 | 30.00 | 0.0062 |
|  | BL vs. Hour 3 | 1.188 | 30.00 | >0.9999 | | BL vs. Hour 3 | 1.782 | 30.00 | 0.4242 |
|  | BL vs. Hour 5 | 2.376 | 30.00 | 0.1203 | | BL vs. Hour 5 | 2.376 | 30.00 | 0.1203 |
|  | BL vs. Hour 24 | 0.5941 | 30.00 | >0.9999 | | BL vs. Hour 24 | 1.188 | 30.00 | >0.9999 |
|  | BL vs. Hour 48 | 1.188 | 30.00 | >0.9999 | | BL vs. Hour 48 | 0.000 | 30.00 | >0.9999 |
| **8A** | BL vs. Hour 1 | 5.171 | 30.00 | <0.0001 | | BL vs. Hour 1 | 4.654 | 30.00 | 0.0003 |
|  | BL vs. Hour 3 | 5.364 | 30.00 | <0.0001 | | BL vs. Hour 3 | 4.889 | 30.00 | 0.0002 |
|  | BL vs. Hour 5 | 5.355 | 30.00 | <0.0001 | | BL vs. Hour 5 | 5.794 | 30.00 | <0.0001 |
|  | BL vs. Hour 24 | 3.499 | 30.00 | 0.0074 | | BL vs. Hour 24 | 3.936 | 30.00 | 0.0023 |
|  | BL vs. Hour 48 | 2.094 | 30.00 | 0.2239 | | BL vs. Hour 48 | 2.165 | 30.00 | 0.1923 |
| **8B** | BL vs. Hour 1 | 3.953 | 30.00 | 0.0022 | | BL vs. Hour 1 | 5.929 | 30.00 | <0.0001 |
|  | BL vs. Hour 3 | 9.882 | 30.00 | <0.0001 | | BL vs. Hour 3 | 9.092 | 30.00 | <0.0001 |
|  | BL vs. Hour 5 | 5.139 | 30.00 | <0.0001 | | BL vs. Hour 5 | 7.115 | 30.00 | <0.0001 |
|  | BL vs. Hour 24 | 5.534 | 30.00 | <0.0001 | | BL vs. Hour 24 | 2.372 | 30.00 | 0.1216 |
|  | BL vs. Hour 48 | 1.581 | 30.00 | 0.6217 | | BL vs. Hour 48 | 1.581 | 30.00 | 0.6217 |
| **8C** | BL vs. Hour 1 | 3.436 | 30.00 | 0.0088 | | BL vs. Hour 1 | 1.055 | 30.00 | >0.9999 |
|  | BL vs. Hour 3 | 5.272 | 30.00 | <0.0001 | | BL vs. Hour 3 | 1.675 | 30.00 | 0.5218 |
|  | BL vs. Hour 5 | 5.705 | 30.00 | <0.0001 | | BL vs. Hour 5 | 0.4832 | 30.00 | >0.9999 |
|  | BL vs. Hour 24 | 5.536 | 30.00 | <0.0001 | | BL vs. Hour 24 | 0.1900 | 30.00 | >0.9999 |
|  | BL vs. Hour 48 | 2.857 | 30.00 | 0.0385 | | BL vs. Hour 48 | 0.4548 | 30.00 | >0.9999 |
| **8D** | BL vs. Hour 1 | 3.415 | 30.00 | 0.0093 | | BL vs. Hour 1 | 0.8537 | 30.00 | >0.9999 |
|  | BL vs. Hour 3 | 8.963 | 30.00 | <0.0001 | | BL vs. Hour 3 | 4.695 | 30.00 | 0.0003 |
|  | BL vs. Hour 5 | 6.829 | 30.00 | <0.0001 | | BL vs. Hour 5 | 2.988 | 30.00 | 0.0278 |
|  | BL vs. Hour 24 | 4.695 | 30.00 | 0.0003 | | BL vs. Hour 24 | 2.561 | 30.00 | 0.0785 |
|  | BL vs. Hour 48 | 2.988 | 30.00 | 0.0278 | | BL vs. Hour 48 | 0.4268 | 30.00 | >0.9999 |

**Supplementary Table 3** – Two-way ANOVA with Bonferroni’s Multiple Comparisons

| **Two-way ANOVA Bonferroni’s Multiple Comparisons** | | | | | | | | | | | | |
| --- | --- | --- | --- | --- | --- | --- | --- | --- | --- | --- | --- | --- |
| Fig | Comparison | (t,DF), p | | | | Fig | Comparison | (t,DF), p | | | | |
|  | WT - PAR3^-/-^ | | | | |  | WT - PAR3^-/-^ | | | | | |
|  |  | t | | DF | p |  |  | t | | DF | p | |
| **S2A** | BL | 0.2148 | | 36.00 | >0.9999 | **S5B** | BL | 0.4529 | | 36.00 | >0.9999 | |
|  | Hour 1 | 0.9446 | | 36.00 | >0.9999 |  | Hour 1 | 0.4529 | | 36.00 | >0.9999 | |
|  | Hour 3 | 0.3352 | | 36.00 | >0.9999 |  | Hour 3 | 0.2265 | | 36.00 | >0.9999 | |
|  | Hour 5 | 0.2611 | | 36.00 | >0.9999 |  | Hour 5 | 0.000 | | 36.00 | >0.9999 | |
|  | Hour 24 | 0.4427 | | 36.00 | >0.9999 |  | Hour 24 | 0.000 | | 36.00 | >0.9999 | |
|  | Hour 48 | 0.2704 | | 36.00 | >0.9999 |  | Hour 48 | 1.359 | | 36.00 | >0.9999 | |
| **S2B** | BL | 0.1034 | | 36.00 | >0.9999 | **S5C** | BL | 0.04047 | | 36.00 | >0.9999 | |
|  | Hour 1 | 0.4135 | | 36.00 | >0.9999 |  | Hour 1 | 0.3821 | | 36.00 | >0.9999 | |
|  | Hour 3 | 1.059 | | 36.00 | >0.9999 |  | Hour 3 | 0.5515 | | 36.00 | >0.9999 | |
|  | Hour 5 | 2.558 | | 36.00 | 0.0892 |  | Hour 5 | 0.7642 | | 36.00 | >0.9999 | |
|  | Hour 24 | 0.02584 | | 36.00 | >0.9999 |  | Hour 24 | 0.6259 | | 36.00 | >0.9999 | |
|  | Hour 48 | 0.5427 | | 36.00 | >0.9999 |  | Hour 48 | 0.000 | | 36.00 | >0.9999 | |
| **S2C** | BL | 0.6108 | | 36.00 | >0.9999 | **S5D** | BL | 0.4027 | | 36.00 | >0.9999 | |
|  | Hour 1 | 2.189 | | 36.00 | 0.2111 |  | Hour 1 | 0.8054 | | 36.00 | >0.9999 | |
|  | Hour 3 | 0.4585 | | 36.00 | >0.9999 |  | Hour 3 | 0.4027 | | 36.00 | >0.9999 | |
|  | Hour 5 | 0.000 | | 36.00 | >0.9999 |  | Hour 5 | 1.208 | | 36.00 | >0.9999 | |
|  | Hour 24 | 0.2120 | | 36.00 | >0.9999 |  | Hour 24 | 0.4027 | | 36.00 | >0.9999 | |
|  | Hour 48 | 0.5935 | | 36.00 | >0.9999 |  | Hour 48 | 0.4027 | | 36.00 | >0.9999 | |
| **S2D** | BL | 0.1764 | | 36.00 | >0.9999 | **S6A** | BL | 0.7152 | | 36.00 | >0.9999 | |
|  | Hour 1 | 2.029 | | 36.00 | 0.2994 |  | Hour 1 | 1.177 | | 36.00 | >0.9999 | |
|  | Hour 3 | 0.7646 | | 36.00 | >0.9999 |  | Hour 3 | 0.04716 | | 36.00 | >0.9999 | |
|  | Hour 5 | 0.05881 | | 36.00 | >0.9999 |  | Hour 5 | 0.6316 | | 36.00 | >0.9999 | |
|  | Hour 24 | 0.1470 | | 36.00 | >0.9999 |  | Hour 24 | 1.274 | | 36.00 | >0.9999 | |
|  | Hour 48 | 0.3529 | | 36.00 | >0.9999 |  | Hour 48 | 0.8173 | | 36.00 | >0.9999 | |
| **S3A** | BL | 0.1175 | | 84.00 | >0.9999 | **S6B** | BL | 0.4588 | | 36.00 | >0.9999 | |
|  | Hour 1 | 1.171 | | 84.00 | >0.9999 |  | Hour 1 | 0.6882 | | 36.00 | >0.9999 | |
|  | Hour 3 | 0.2222 | | 84.00 | >0.9999 |  | Hour 3 | 0.9177 | | 36.00 | >0.9999 | |
|  | Hour 5 | 0.4422 | | 84.00 | >0.9999 |  | Hour 5 | 3.212 | | 36.00 | 0.0167 | |
|  | Hour 24 | 1.590 | | 84.00 | 0.6933 |  | Hour 24 | 1.147 | | 36.00 | >0.9999 | |
|  | Hour 48 | 0.5220 | | 84.00 | >0.9999 |  | Hour 48 | 1.376 | | 36.00 | >0.9999 | |
| **S3B** | BL | 0.4657 | | 36.00 | >0.9999 | **S6C** | BL | 0.2635 | | 36.00 | >0.9999 | |
|  | Hour 1 | 1.397 | | 36.00 | >0.9999 |  | Hour 1 | 5.482 | | 36.00 | <0.0001 | |
|  | Hour 3 | 0.4657 | | 36.00 | >0.9999 |  | Hour 3 | 5.516 | | 36.00 | <0.0001 | |
|  | Hour 5 | 0.4657 | | 36.00 | >0.9999 |  | Hour 5 | 7.464 | | 36.00 | <0.0001 | |
|  | Hour 24 | 0.9314 | | 36.00 | >0.9999 |  | Hour 24 | 6.184 | | 36.00 | <0.0001 | |
|  | Hour 48 | 0.4657 | | 36.00 | >0.9999 |  | Hour 48 | 1.180 | | 36.00 | >0.9999 | |
| **S3C** | BL | 0.5914 | | 84.00 | >0.9999 | **S6D** | BL | 0.000 | | 36.00 | >0.9999 | |
|  | Hour 1 | 0.2041 | | 84.00 | >0.9999 |  | Hour 1 | 0.000 | | 36.00 | >0.9999 | |
|  | Hour 3 | 0.2532 | | 84.00 | >0.9999 |  | Hour 3 | 4.346 | | 36.00 | 0.0007 | |
|  | Hour 5 | 1.547 | | 84.00 | 0.7543 |  | Hour 5 | 3.259 | | 36.00 | 0.0147 | |
|  | Hour 24 | 0.6642 | | 84.00 | >0.9999 |  | Hour 24 | 3.803 | | 36.00 | 0.0032 | |
|  | Hour 48 | 0.8726 | | 84.00 | >0.9999 |  | Hour 48 | 2.173 | | 36.00 | 0.2187 | |
| **S3D** | BL | 0.3873 | | 36.00 | >0.9999 | **S7A** | BL | 0.00855 | | 36.00 | >0.9999 | |
|  | Hour 1 | 1.162 | | 36.00 | >0.9999 |  | Hour 1 | 0.2482 | | 36.00 | >0.9999 | |
|  | Hour 3 | 0.7746 | | 36.00 | >0.9999 |  | Hour 3 | 0.2274 | | 36.00 | >0.9999 | |
|  | Hour 5 | 0.7746 | | 36.00 | >0.9999 |  | Hour 5 | 1.380 | | 36.00 | >0.9999 | |
|  | Hour 24 | 1.549 | | 36.00 | 0.7805 |  | Hour 24 | 0.2678 | | 36.00 | >0.9999 | |
|  | Hour 48 | 0.7746 | | 36.00 | >0.9999 |  | Hour 48 | 0.8665 | | 36.00 | >0.9999 | |
| **S4A** | BL | 0.3954 | | 36.00 | >0.9999 | **S7B** | BL | 0.3762 | | 36.00 | >0.9999 | |
|  | Hour 1 | 2.401 | | 36.00 | 0.1299 |  | Hour 1 | 0.7524 | | 36.00 | >0.9999 | |
|  | Hour 3 | 5.780 | | 36.00 | <0.0001 |  | Hour 3 | 0.3762 | | 36.00 | >0.9999 | |
|  | Hour 5 | 0.2470 | | 36.00 | >0.9999 |  | Hour 5 | 1.129 | | 36.00 | >0.9999 | |
|  | Hour 24 | 1.768 | | 36.00 | 0.5135 |  | Hour 24 | 0.4702 | | 36.00 | >0.9999 | |
|  | Hour 48 | 0.9633 | | 36.00 | >0.9999 |  | Hour 48 | 8.353e-016 | | 36.00 | >0.9999 | |
| **S4B** | BL | 0.000 | | 36.00 | >0.9999 | **S7C** | BL | 0.000 | | 36.00 | >0.9999 | |
|  | Hour 1 | 2.631 | | 36.00 | 0.0747 |  | Hour 1 | 1.346 | | 36.00 | >0.9999 | |
|  | Hour 3 | 1.579 | | 36.00 | 0.7389 |  | Hour 3 | 0.000 | | 36.00 | >0.9999 | |
|  | Hour 5 | 0.5262 | | 36.00 | >0.9999 |  | Hour 5 | 4.447 | | 36.00 | 0.0005 | |
|  | Hour 24 | 0.000 | | 36.00 | >0.9999 |  | Hour 24 | 5.567 | | 36.00 | <0.0001 | |
|  | Hour 48 | 2.105 | | 36.00 | 0.2540 |  | Hour 48 | 2.148 | | 36.00 | 0.2313 | |
| **S5A** | BL | 0.3524 | | 36.00 | >0.9999 | **S7D** | BL | 0.5222 | | 36.00 | >0.9999 | |
|  | Hour 1 | 1.205 | | 36.00 | >0.9999 |  | Hour 1 | 0.000 | | 36.00 | >0.9999 | |
|  | Hour 3 | 1.318 | | 36.00 | >0.9999 |  | Hour 3 | 0.5222 | | 36.00 | >0.9999 | |
|  | Hour 5 | 0.3880 | | 36.00 | >0.9999 |  | Hour 5 | 4.178 | | 36.00 | 0.0011 | |
|  | Hour 24 | 0.2688 | | 36.00 | >0.9999 |  | Hour 24 | 1.044 | | 36.00 | >0.9999 | |
|  | Hour 48 | 0.9051 | | 36.00 | >0.9999 |  | Hour 48 | 2.089 | | 36.00 | 0.2631 | |
| **Unpaired t-test** | | | | | | | | | | | | |
| Fig | (t,DF), p | | | | | Fig | (t,DF), p | | | | | |
| **S2A** | 0.4264 | | 6 | 0.6847 | | **S5B** | 0.6449 | | 6 | | | 0.5428 |
| **S2B** | 0.8530 | | 6 | 0.4264 | | **S5C** | 0.2231 | | 6 | | | 0.8309 |
| **S2C** | 0.8142 | | 6 | 0.4467 | | **S5D** | 0.09298 | | 6 | | | 0.9289 |
| **S2D** | 0.4174 | | 6 | 0.6909 | | **S6A** | 2.044 | | 6 | | | 0.0869 |
| **S3A** | 0.5042 | | 14 | 0.6220 | | **S6B** | 1.934 | | 6 | | | 0.1012 |
| **S3B** | 0.4229 | | 6 | 0.6871 | | **S6C** | 5.420 | | 6 | | | 0.0016 |
| **S3C** | 1.217 | | 14 | 0.2436 | | **S6D** | 2.488 | | 6 | | | 0.0473 |
| **S3D** | 2.274 | | 6 | 0.0633 | | **S7A** | 0.3660 | | 6 | | | 0.7269 |
| **S4A** | 1.859 | | 6 | 0.1124 | | **S7B** | 0.5610 | | 6 | | | 0.5951 |
| **S4B** | 0.09637 | | 6 | 0.9264 | | **S7C** | 3.856 | | 6 | | | 0.0084 |
| **S5A** | 0.8996 | | 6 | 0.4030 | | **S7D** | 2.056 | | 6 | | | 0.0856 |

**Supplementary Table 4** - Two-way ANOVA with Bonferroni’s Multiple Comparisons

| Fig | Comparison | (t,DF), p | | | Comparison | (t,DF), p | | |
| --- | --- | --- | --- | --- | --- | --- | --- | --- |
|  | WT | | | | PAR3^-/-^ | | | |
|  |  | t | DF | p |  | t | DF | p |
| **S2AAAAA** | BL vs. Hour 1 | 0.1219 | 30.00 | >0.9999 | BL vs. Hour 1 | 0.8385 | 30.00 | >0.9999 |
|  | BL vs. Hour 3 | 0.9822 | 30.00 | >0.9999 | BL vs. Hour 3 | 1.522 | 30.00 | 0.6919 |
|  | BL vs. Hour 5 | 1.357 | 30.00 | 0.9247 | BL vs. Hour 5 | 0.8894 | 30.00 | >0.9999 |
|  | BL vs. Hour 24 | 1.228 | 30.00 | >0.9999 | BL vs. Hour 24 | 0.5820 | 30.00 | >0.9999 |
|  | BL vs. Hour 48 | 0.8167 | 30.00 | >0.9999 | BL vs. Hour 48 | 0.8712 | 30.00 | >0.9999 |
| **S2B** | BL vs. Hour 1 | 0.9382 | 30.00 | >0.9999 | BL vs. Hour 1 | 0.6071 | 30.00 | >0.9999 |
|  | BL vs. Hour 3 | 0.3311 | 30.00 | >0.9999 | BL vs. Hour 3 | 0.9106 | 30.00 | >0.9999 |
|  | BL vs. Hour 5 | 2.594 | 30.00 | 0.0727 | BL vs. Hour 5 | 0.2483 | 30.00 | >0.9999 |
|  | BL vs. Hour 24 | 0.3311 | 30.00 | >0.9999 | BL vs. Hour 24 | 0.1932 | 30.00 | >0.9999 |
|  | BL vs. Hour 48 | 0.8278 | 30.00 | >0.9999 | BL vs. Hour 48 | 0.1380 | 30.00 | >0.9999 |
| **S2C** | BL vs. Hour 1 | 3.237 | 30.00 | 0.0147 | BL vs. Hour 1 | 0.6077 | 30.00 | >0.9999 |
|  | BL vs. Hour 3 | 0.2065 | 30.00 | >0.9999 | BL vs. Hour 3 | 0.7979 | 30.00 | >0.9999 |
|  | BL vs. Hour 5 | 1.599 | 30.00 | 0.6017 | BL vs. Hour 5 | 1.025 | 30.00 | >0.9999 |
|  | BL vs. Hour 24 | 1.229 | 30.00 | >0.9999 | BL vs. Hour 24 | 0.4558 | 30.00 | >0.9999 |
|  | BL vs. Hour 48 | 0.2935 | 30.00 | >0.9999 | BL vs. Hour 48 | 0.2773 | 30.00 | >0.9999 |
| **S2D** | BL vs. Hour 1 | 0.4852 | 30.00 | >0.9999 | BL vs. Hour 1 | 1.425 | 30.00 | 0.8219 |
|  | BL vs. Hour 3 | 0.4246 | 30.00 | >0.9999 | BL vs. Hour 3 | 0.1820 | 30.00 | >0.9999 |
|  | BL vs. Hour 5 | 0.9098 | 30.00 | >0.9999 | BL vs. Hour 5 | 0.6672 | 30.00 | >0.9999 |
|  | BL vs. Hour 24 | 0.09098 | 30.00 | >0.9999 | BL vs. Hour 24 | 0.4246 | 30.00 | >0.9999 |
|  | BL vs. Hour 48 | 0.1516 | 30.00 | >0.9999 | BL vs. Hour 48 | 0.3942 | 30.00 | >0.9999 |
| **S3A** | BL vs. Hour 1 | 2.458 | 70.00 | 0.0822 | BL vs. Hour 1 | 3.490 | 70.00 | 0.0042 |
|  | BL vs. Hour 3 | 8.442 | 70.00 | <0.0001 | BL vs. Hour 3 | 8.545 | 70.00 | <0.0001 |
|  | BL vs. Hour 5 | 11.37 | 70.00 | <0.0001 | BL vs. Hour 5 | 10.83 | 70.00 | <0.0001 |
|  | BL vs. Hour 24 | 5.551 | 70.00 | <0.0001 | BL vs. Hour 24 | 3.879 | 70.00 | 0.0012 |
|  | BL vs. Hour 48 | 1.257 | 70.00 | >0.9999 | BL vs. Hour 48 | 0.6305 | 70.00 | >0.9999 |
| **S3B** | BL vs. Hour 1 | 4.363 | 30.00 | 0.0007 | BL vs. Hour 1 | 5.332 | 30.00 | <0.0001 |
|  | BL vs. Hour 3 | 12.12 | 30.00 | <0.0001 | BL vs. Hour 3 | 11.15 | 30.00 | <0.0001 |
|  | BL vs. Hour 5 | 10.18 | 30.00 | <0.0001 | BL vs. Hour 5 | 10.18 | 30.00 | <0.0001 |
|  | BL vs. Hour 24 | 6.302 | 30.00 | <0.0001 | BL vs. Hour 24 | 4.848 | 30.00 | 0.0002 |
|  | BL vs. Hour 48 | 0.9695 | 30.00 | >0.9999 | BL vs. Hour 48 | 0.9695 | 30.00 | >0.9999 |
| **S3C** | BL vs. Hour 1 | 7.935 | 70.00 | <0.0001 | BL vs. Hour 1 | 8.324 | 70.00 | <0.0001 |
|  | BL vs. Hour 3 | 8.895 | 70.00 | <0.0001 | BL vs. Hour 3 | 9.235 | 70.00 | <0.0001 |
|  | BL vs. Hour 5 | 8.288 | 70.00 | <0.0001 | BL vs. Hour 5 | 10.44 | 70.00 | <0.0001 |
|  | BL vs. Hour 24 | 2.948 | 70.00 | 0.0217 | BL vs. Hour 24 | 4.211 | 70.00 | 0.0004 |
|  | BL vs. Hour 48 | 0.6220 | 70.00 | >0.9999 | BL vs. Hour 48 | 0.8501 | 70.00 | >0.9999 |
| **S3D** | BL vs. Hour 1 | 4.394 | 30.00 | 0.0006 | BL vs. Hour 1 | 5.193 | 30.00 | <0.0001 |
|  | BL vs. Hour 3 | 9.188 | 30.00 | <0.0001 | BL vs. Hour 3 | 7.989 | 30.00 | <0.0001 |
|  | BL vs. Hour 5 | 5.593 | 30.00 | <0.0001 | BL vs. Hour 5 | 4.394 | 30.00 | 0.0006 |
|  | BL vs. Hour 24 | 1.598 | 30.00 | 0.6028 | BL vs. Hour 24 | 2.796 | 30.00 | 0.0447 |
|  | BL vs. Hour 48 | 1.598 | 30.00 | 0.6028 | BL vs. Hour 48 | 0.3995 | 30.00 | >0.9999 |
| **S4A** | BL vs. Hour 1 | 4.354 | 30.00 | 0.0007 | BL vs. Hour 1 | 2.331 | 30.00 | 0.1332 |
|  | BL vs. Hour 3 | 7.693 | 30.00 | <0.0001 | BL vs. Hour 3 | 2.262 | 30.00 | 0.1555 |
|  | BL vs. Hour 5 | 2.935 | 30.00 | 0.0317 | BL vs. Hour 5 | 3.085 | 30.00 | 0.0218 |
|  | BL vs. Hour 24 | 3.945 | 30.00 | 0.0022 | BL vs. Hour 24 | 2.561 | 30.00 | 0.0786 |
|  | BL vs. Hour 48 | 0.4372 | 30.00 | >0.9999 | BL vs. Hour 48 | 0.9331 | 30.00 | >0.9999 |
| **S4B** | BL vs. Hour 1 | 4.034 | 30.00 | 0.0017 | BL vs. Hour 1 | 1.153 | 30.00 | >0.9999 |
|  | BL vs. Hour 3 | 2.881 | 30.00 | 0.0362 | BL vs. Hour 3 | 1.153 | 30.00 | >0.9999 |
|  | BL vs. Hour 5 | 0.5763 | 30.00 | >0.9999 | BL vs. Hour 5 | 1.153 | 30.00 | >0.9999 |
|  | BL vs. Hour 24 | 2.305 | 30.00 | 0.1413 | BL vs. Hour 24 | 2.305 | 30.00 | 0.1413 |
|  | BL vs. Hour 48 | 2.305 | 30.00 | 0.1413 | BL vs. Hour 48 | 0.000 | 30.00 | >0.9999 |
| **S5A** | BL vs. Hour 1 | 3.852 | 30.00 | 0.0029 | BL vs. Hour 1 | 2.996 | 30.00 | 0.0273 |
|  | BL vs. Hour 3 | 6.039 | 30.00 | <0.0001 | BL vs. Hour 3 | 7.717 | 30.00 | <0.0001 |
|  | BL vs. Hour 5 | 6.246 | 30.00 | <0.0001 | BL vs. Hour 5 | 6.990 | 30.00 | <0.0001 |
|  | BL vs. Hour 24 | 4.122 | 30.00 | 0.0014 | BL vs. Hour 24 | 4.206 | 30.00 | 0.0011 |
|  | BL vs. Hour 48 | 0.5553 | 30.00 | >0.9999 | BL vs. Hour 48 | 1.819 | 30.00 | 0.3946 |
| **S5B** | BL vs. Hour 1 | 2.599 | 30.00 | 0.0718 | BL vs. Hour 1 | 2.599 | 30.00 | 0.0718 |
|  | BL vs. Hour 3 | 11.91 | 30.00 | <0.0001 | BL vs. Hour 3 | 12.13 | 30.00 | <0.0001 |
|  | BL vs. Hour 5 | 10.83 | 30.00 | <0.0001 | BL vs. Hour 5 | 11.26 | 30.00 | <0.0001 |
|  | BL vs. Hour 24 | 2.166 | 30.00 | 0.1921 | BL vs. Hour 24 | 2.599 | 30.00 | 0.0718 |
|  | BL vs. Hour 48 | 9.617e-016 | 30.00 | >0.9999 | BL vs. Hour 48 | 1.733 | 30.00 | 0.4673 |
| **S5C** | BL vs. Hour 1 | 6.826 | 30.00 | <0.0001 | BL vs. Hour 1 | 7.285 | 30.00 | <0.0001 |
|  | BL vs. Hour 3 | 10.77 | 30.00 | <0.0001 | BL vs. Hour 3 | 11.41 | 30.00 | <0.0001 |
|  | BL vs. Hour 5 | 10.39 | 30.00 | <0.0001 | BL vs. Hour 5 | 9.600 | 30.00 | <0.0001 |
|  | BL vs. Hour 24 | 3.840 | 30.00 | 0.0030 | BL vs. Hour 24 | 4.563 | 30.00 | 0.0004 |
|  | BL vs. Hour 48 | 0.8960 | 30.00 | >0.9999 | BL vs. Hour 48 | 0.9399 | 30.00 | >0.9999 |
| **S5D** | BL vs. Hour 1 | 2.098 | 30.00 | 0.2220 | BL vs. Hour 1 | 2.518 | 30.00 | 0.0869 |
|  | BL vs. Hour 3 | 7.134 | 30.00 | <0.0001 | BL vs. Hour 3 | 6.295 | 30.00 | <0.0001 |
|  | BL vs. Hour 5 | 6.715 | 30.00 | <0.0001 | BL vs. Hour 5 | 7.554 | 30.00 | <0.0001 |
|  | BL vs. Hour 24 | 0.8393 | 30.00 | >0.9999 | BL vs. Hour 24 | 0.8393 | 30.00 | >0.9999 |
|  | BL vs. Hour 48 | 0.4197 | 30.00 | >0.9999 | BL vs. Hour 48 | 0.4197 | 30.00 | >0.9999 |
| **S6A** | BL vs. Hour 1 | 5.955 | 30.00 | <0.0001 | BL vs. Hour 1 | 4.067 | 30.00 | 0.0016 |
|  | BL vs. Hour 3 | 4.434 | 30.00 | 0.0006 | BL vs. Hour 3 | 3.767 | 30.00 | 0.0036 |
|  | BL vs. Hour 5 | 4.663 | 30.00 | 0.0003 | BL vs. Hour 5 | 3.320 | 30.00 | 0.0119 |
|  | BL vs. Hour 24 | 4.551 | 30.00 | 0.0004 | BL vs. Hour 24 | 2.567 | 30.00 | 0.0774 |
|  | BL vs. Hour 48 | 3.346 | 30.00 | 0.0111 | BL vs. Hour 48 | 1.817 | 30.00 | 0.3957 |
| **S6B** | BL vs. Hour 1 | 2.791 | 30.00 | 0.0453 | BL vs. Hour 1 | 2.558 | 30.00 | 0.0791 |
|  | BL vs. Hour 3 | 2.093 | 30.00 | 0.2245 | BL vs. Hour 3 | 1.628 | 30.00 | 0.5700 |
|  | BL vs. Hour 5 | 3.256 | 30.00 | 0.0140 | BL vs. Hour 5 | 0.4651 | 30.00 | >0.9999 |
|  | BL vs. Hour 24 | 1.395 | 30.00 | 0.8657 | BL vs. Hour 24 | 0.6977 | 30.00 | >0.9999 |
|  | BL vs. Hour 48 | 1.163 | 30.00 | >0.9999 | BL vs. Hour 48 | 0.2326 | 30.00 | >0.9999 |
| **S6C** | BL vs. Hour 1 | 6.748 | 30.00 | <0.0001 | BL vs. Hour 1 | 1.474 | 30.00 | 0.7546 |
|  | BL vs. Hour 3 | 7.363 | 30.00 | <0.0001 | BL vs. Hour 3 | 2.054 | 30.00 | 0.2437 |
|  | BL vs. Hour 5 | 9.576 | 30.00 | <0.0001 | BL vs. Hour 5 | 2.298 | 30.00 | 0.1436 |
|  | BL vs. Hour 24 | 6.924 | 30.00 | <0.0001 | BL vs. Hour 24 | 0.9400 | 30.00 | >0.9999 |
|  | BL vs. Hour 48 | 1.675 | 30.00 | 0.5220 | BL vs. Hour 48 | 0.7482 | 30.00 | >0.9999 |
| **S6D** | BL vs. Hour 1 | 10.49 | 30.00 | <0.0001 | BL vs. Hour 1 | 10.49 | 30.00 | <0.0001 |
|  | BL vs. Hour 3 | 7.869 | 30.00 | <0.0001 | BL vs. Hour 3 | 3.672 | 30.00 | 0.0047 |
|  | BL vs. Hour 5 | 5.246 | 30.00 | <0.0001 | BL vs. Hour 5 | 2.098 | 30.00 | 0.2219 |
|  | BL vs. Hour 24 | 4.722 | 30.00 | 0.0003 | BL vs. Hour 24 | 1.049 | 30.00 | >0.9999 |
|  | BL vs. Hour 48 | 2.098 | 30.00 | 0.2219 | BL vs. Hour 48 | 5.824e-016 | 30.00 | >0.9999 |
| **S7A** | BL vs. Hour 1 | 3.644 | 30.00 | 0.0050 | BL vs. Hour 1 | 3.892 | 30.00 | 0.0026 |
|  | BL vs. Hour 3 | 7.777 | 30.00 | <0.0001 | BL vs. Hour 3 | 8.003 | 30.00 | <0.0001 |
|  | BL vs. Hour 5 | 8.012 | 30.00 | <0.0001 | BL vs. Hour 5 | 6.573 | 30.00 | <0.0001 |
|  | BL vs. Hour 24 | 6.267 | 30.00 | <0.0001 | BL vs. Hour 24 | 6.535 | 30.00 | <0.0001 |
|  | BL vs. Hour 48 | 4.368 | 30.00 | 0.0007 | BL vs. Hour 48 | 3.462 | 30.00 | 0.0082 |
| **S7B** | BL vs. Hour 1 | 1.642 | 30.00 | 0.5549 | BL vs. Hour 1 | 2.874 | 30.00 | 0.0369 |
|  | BL vs. Hour 3 | 6.980 | 30.00 | <0.0001 | BL vs. Hour 3 | 6.980 | 30.00 | <0.0001 |
|  | BL vs. Hour 5 | 4.927 | 30.00 | 0.0001 | BL vs. Hour 5 | 4.106 | 30.00 | 0.0014 |
|  | BL vs. Hour 24 | 1.232 | 30.00 | >0.9999 | BL vs. Hour 24 | 2.155 | 30.00 | 0.1964 |
|  | BL vs. Hour 48 | 4.558e-016 | 30.00 | >0.9999 | BL vs. Hour 48 | 0.4106 | 30.00 | >0.9999 |
| **S7C** | BL vs. Hour 1 | 2.576 | 30.00 | 0.0759 | BL vs. Hour 1 | 1.326 | 30.00 | 0.9741 |
|  | BL vs. Hour 3 | 3.287 | 30.00 | 0.0129 | BL vs. Hour 3 | 3.287 | 30.00 | 0.0129 |
|  | BL vs. Hour 5 | 6.933 | 30.00 | <0.0001 | BL vs. Hour 5 | 2.803 | 30.00 | 0.0440 |
|  | BL vs. Hour 24 | 6.000 | 30.00 | <0.0001 | BL vs. Hour 24 | 0.8298 | 30.00 | >0.9999 |
|  | BL vs. Hour 48 | 2.295 | 30.00 | 0.1446 | BL vs. Hour 48 | 0.3004 | 30.00 | >0.9999 |
| **S7D** | BL vs. Hour 1 | 2.460 | 30.00 | 0.0994 | BL vs. Hour 1 | 1.845 | 30.00 | 0.3748 |
|  | BL vs. Hour 3 | 1.845 | 30.00 | 0.3748 | BL vs. Hour 3 | 1.845 | 30.00 | 0.3748 |
|  | BL vs. Hour 5 | 5.534 | 30.00 | <0.0001 | BL vs. Hour 5 | 0.000 | 30.00 | >0.9999 |
|  | BL vs. Hour 24 | 1.845 | 30.00 | 0.3748 | BL vs. Hour 24 | 0.000 | 30.00 | >0.9999 |
|  | BL vs. Hour 48 | 2.460 | 30.00 | 0.0994 | BL vs. Hour 48 | 0.6149 | 30.00 | >0.9999 |
